# High-pH NMR to Identify Macromolecular Hydrogen-Bonds and Foldons

**DOI:** 10.64898/2026.02.28.708709

**Authors:** Andrei T. Alexandrescu, Antonio J. Rua, Sameena Shah, Daniel Fairchild, Irina Bezsonova

## Abstract

Hydrogen bond (H-bond) restraints are critical for NMR structure determination, yet their experimental identification can be challenging for marginally stable structures that afford insufficient protection from (H/D) exchange in D_2_O. As an alternative, we explored the use of NMR between pH 10 and 11 conditions that promote rapid exchange, for identifying backbone amide protons involved in H-bonds. We analyzed ∼750 amide sites distributed across ten proteins with known structures. Persistence of amide protons at high pH in standard 2D ^1^H-^15^N HSQC spectra for ^15^N-labeled proteins in H2O, or TOCSY for unlabeled proteins, identifies H-bonds with ∼91% accuracy that exceeds the ∼80% accuracy of traditional H/D exchange experiments in D_2_O. For two α-helical coiled coils and three globular proteins, we performed alkaline unfolding experiments taking advantage of amide NMR signal attenuation from unstructured polypeptides. Increasing the sample pH led to a progressive loss of native amide proton NMR signals, revealing an unfolding hierarchy where “foldons” remaining at the highest pH values had the most persistent H-bonds under EX1 exchange conditions. The foldons observed at high pH are consistent with partially folded structures previously characterized near neutral pH by native state hydrogen exchange, equilibrium unfolding, and protein fragment studies. For β-sheet proteins, foldons correspond to regions with high inter-residue contact density, whereas in coiled coils they demarcate regions with high α-helical propensity. High-pH NMR experiments provide a sensitive, fast, inexpensive, and broadly applicable approach to map H-bonding in marginally stable or partially folded proteins. Additionally, they offer the opportunity to explore uncharted protein dynamics and unfolding pathways under basic pH conditions.

## 1. INTRODUCTION

Hydrogen bonds (H-bonds) are cornerstones of protein structures contributing to their stabilities and folding specificities, as well as forming regular interaction patterns such as α-helix and β-sheet secondary structures [1–3]. In NMR structure determination, H-bond restraints greatly improve conformational convergence even when supplemented with only sparse experimental data [4, 5]. By contrast, without explicit H-bond constraints NMR structures rarely approach the accuracy of the gold-standard X-ray structures, or even of Alpha Fold models [6].

Historically, H-bonds were included in NMR structure calculations only when validated by hydrogen–deuterium (H/D) isotope exchange in D₂O, where amide proton protection from solvent was inferred to indicate H-bonding [7]. This protection requirement is overly stringent as exchange depends on both H-bonding and stability [8–10], with weak or transient H-bonds failing to afford exchange protection. Computational approaches such as ANSURR infer and iteratively include H-bonds during cycles of NMR structure refinement but ultimately depend on validation against experimentally determined H-bond datasets [5]. H-bonds can be detected directly by NMR using long-range HNCO experiments that establish through-H-bond ^h³^J_NC′_ couplings. These experiments identify both the H-bond donor and acceptor [11, 12] but depend on the measurement of very small 0.3-1 Hz couplings that require long delays (∼133 ms) to detect, and therefore typically need perdeuterated samples for molecular weights above 5-10 KDa [13]. These factors combine to make the use of long-range HNCO experiments for determining H-bonds for NMR structure determination rare, with most studies still relying on H/D exchange in D_2_O to identify H-bonded sites. Because H/D exchange depends on structural stability as well as environmental variables such as temperature and pH, protection can be undetectable even close to optimal conditions: low temperature and near the ∼ pH 4.5 acidic minimum for exchange [14]. Examples include the unstable SN-OB fragment of staphylococcal nuclease [10], the small protein CspA in the absence of stabilizing osmolytes [15], as well as several small zinc fingers that are a recent interest of our lab [16–18]. Particularly problematic are partially folded proteins and folding intermediates where H-bonds rarely persist long enough to afford detectable protection [19].

To overcome the limitations of H/D isotope exchange that amide protons need to survive for at least a few minutes to be detected, we examined whether high-pH NMR could serve as an alternative method to identify H-bonds. Since HX is base-catalyzed, rates increase 10-fold with every pH unit above the pH 4.5 minimum for exchange [20]. For structured H-bonded sites, the mechanism of exchange shifts from the EX2 limit, where protection depends on the stability of the H-bonded structure, towards the EX1 limit where protection depends on the rates of H-bonds breaking at high pH [20]. However, even approaching the EX1 limit at pH 10-11, H-bonding will confer amide exchange protection compared to solvent exposed sites. Using a set of ∼750 amide proton sites from ten proteins we established that survival of amide protons identifies H-bonds with high >90% accuracy at the pH 10-11 conditions that promote rapid hydrogen exchange. Moreover, we find that in three globular protein examples and two coiled coils, NMR data at progressively higher pH, reveal a hierarchy of structural stabilities prior to complete alkaline unfolding. This unfolding hierarchy agrees with that established for the proteins by other methods such as equilibrium denaturation, mutagenesis, and native-state HX analyses. Because the high-pH experiments employ sensitive and routine NMR experiments such as ^1^H-^15^N HSQC (or TOCSY for unlabeled samples) they are inexpensive, fast, straightforward, and broadly applicable to probe hydrogen bonding. High-pH NMR offers the opportunity to distinguish folded (protected) and unfolded (unprotected) segments in proteins containing a mixture of structured and disorder regions, and to access largely unexplored information on molecular dynamics and unfolding under alkaline conditions.

## 2. RESULTS

### 2.1 High pH causes loss of amide proton NMR signals in H_2_O due to fast solvent exchange

In typical H/D isotope exchange experiments a lyophilized protein sample is dissolved in D_2_O. Over time, since ^1^H NMR experiments do not detect ^2^H, amide proton NMR signals disappear as the exchange-labile hydrogens on the protein are replaced with deuterons from the solvent [21, 22]. Isotope exchange is effectively irreversible, since ^1^H isotopes from protein concentrations typically in the sub-millimolar range are diluted into a pool of 110 molar D atoms from the D_2_O solvent.

Amide proton NMR signals can also be lost for proteins in H_2_O through a variety of mechanisms. Fig. 1A shows ^1^H-^15^N HSQC spectra of the unstructured Alzheimer’s-associated Aβ(40) peptide in H_2_O at increasing pH values. At pH values below 6.7 (as far down as pH 2.0), all of the 39 expected amide proton crosspeaks from the sequence are observed at a temperature of 20 °C (Fig. 1). The first residue has an α-amine nitrogen and is not observed. As the pH is raised by one pH unit from 6.7 to 7.7, most of the amide protons from G, S and T residues, with ^15^N chemical shifts below 120 ppm in the ^1^H-^15^N HSQC spectrum, are lost. A further increase to pH 8.4 causes the loss of all the signals except for that of the C-terminus V40 (pink arrow), which has the slowest intrinsic exchange (*k_int_*) rate predicted from the amino acid sequence [23]. The results exemplify that the majority of amide protons signals in unstructured proteins and peptides typically disappear over a narrow range of about 1.5 pH units near the physiological pH of 7.4, as seen in numerous other examples such as the intrinsically disordered protein α-synuclein or small unstructured peptides [14, 24]. In fact, protein NMR experiments are typically done at a slightly acidic pH in the range 6.0-6.8 and/or low temperatures, precisely to avoid loss of amide protons through fast exchange [25, 26]. Although amide protons at high pH are usually only lost from unstructured regions, these segments can be crucial for obtaining complete sequential NMR assignments, a requirement for comprehensive NMR studies.

**Figure 1.**
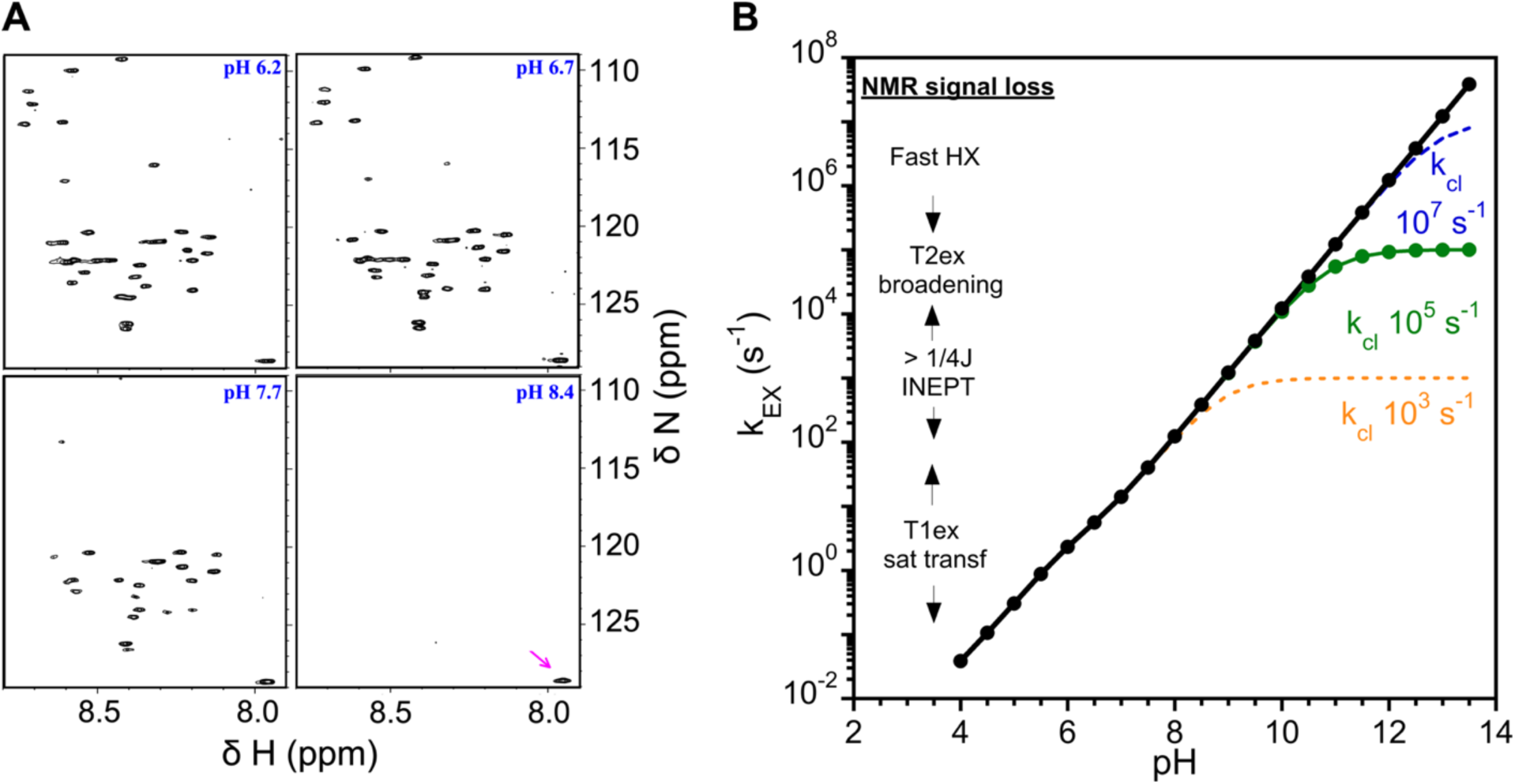
Protein hydrogen exchange in the EX1 limit. **(A)** Selected spectra from a pH titration of ^15^N-labeled Alzheimer’s Aβ(40) peptide at 20 °C. With increasing pH, amide protons from the unstructured Aβ monomers disappear in H_2_O due to increasingly accelerated k_int_ hydrogen exchange rates. **(B)** Intrinsic exchange rates (k_int_) for the Aβ(40) at various pH values in H_2_O at a temperature of 20 °C, predicted from the amino acid sequence with the program SPHERE [23]. Each point at the given pH represents the k_int_ value averaged over the peptide sequence (excluding the N- and C-term residues). Note that the y-axis (k_EX_) is logarithmic so that *k_int_* increase 10-fold with every pH unit. On the left side of the plot are shown k_EX_ ranges for the disappearance of amide proton NMR signals by the various mechanisms described in the main text. The Linderstrom-Lang equation for HX (Eq. 1) was used to simulate the effects of various closing rates (k_cl_) on HX using the k_int_ data for Aβ(40). In all these simulations k_cl_ = k_op_, and therefore K_op_ = 1. The simulations show that the closing rate k_cl_ determines the pH value at which the rate plateau corresponding to the EX1 limit of HX is reached. For a typical protein folding rate (k_cl_ = 10 µs, green curve) the EX1 plateau is reached at about pH 10.5. We did additional simulations keeping k_cl_ constant and varying k_op_ (Fig. S1).

Fig. 1B shows the pH dependence of *k_int_* rates calculated for the Aβ(40) amino acid sequence with the program SPHERE (https://spin.niddk.nih.gov/bax/nmrserver/sphere/ accessed on Dec 21, 2025; [23]). Above the minimum for exchange near pH 4, the intrinsic exchange rate (dependent only on amino acid sequence, pH, and temperature) is expected to increase 10-fold with every unit increase in pH; therefore, a linear relationship is expected between log(*k_int_*) and pH. In the left-side inset for Fig. 1B, ranges of *k_EX_* values are matched with different types of mechanisms for loss of amide proton NMR signals in H_2_O. On the slowest timescale of about 0.1-10 s, amide proton NMR signals can be attenuated through transfer of solvent suppression. Under these conditions, NMR signals from H_2_O and labile protein amide protons are in slow exchange on the chemical shift timescale but in fast exchange on the T1 relaxation timescale.

During the T1 relaxation time, the necessary suppression of the H_2_O signal is partially transferred to exchangeable hydrogens on the protein. The effect is most pronounced using PRESAT type pulse sequences [27], but even more selective solvent suppression schemes, such as those using gradient dephasing, can perturb signals on the protein through exchange, albeit to a reduced extent [28]. Next, timescales faster than 10 ms will cause loss of NMR signal because the amide proton residence time on the protein becomes shorter than the INEPT period (typically (1/4J_HN_) = 2.5 ms) used to establish coupling to the nitrogen. Experiments that use cross-polarization (CP) rather than INEPT to transfer magnetization from ^1^H to ^15^N can circumvent some of these problems and detect faster exchanging amide protons [29]. In another mechanism, amide proton exchange with timescales between about 1 ms and 1µs will cause loss of signal due to T2-exchange broadening contributions. Finally, on the fastest timescales shorter than 1 µs, assuming a typical chemical shift difference of 1-3 ppm or 500-3000 Hz between protein amide and solvent protons on 500 to 1000 MHz instruments, the protein amide and solvent protons will be in fast exchange on the chemical shift timescale. The population-weighted average chemical shift in this case will be dominated by the large excess of solvent compared to the protein and tend towards the ∼4.7 ppm shift of H_2_O. Taken together, these effects combine to cause a progressive loss of amide proton signals in standard ^1^H-^15^N HSQC experiments as the pH is raised beyond about pH 7 [24]. Most amide protons in the Aβ(40) example disappear by about pH 8.5 (Fig. 1A), corresponding to *k_EX_* < 10^3^ s^-1^ (lifetime < 1 ms). This suggests that T2_ex_ broadening and lifetimes shorter than the INEPT coupling period are the dominant mechanisms of NMR signal loss under these conditions (Fig. 1B).

Solvent exchange can be slowed down considerably by protein structure, particularly H-bonded structure. The Linderstorm-Lang model of HX in structured proteins presupposes a *Closed* H-bonded state that is exchange-resistant and an *Open* exchange-susceptible conformation. In this model HX occurs at the *k_int_* rate only from the *Ope*n state, when it is generated through conformational fluctuations from the *Closed* state [30, 31].

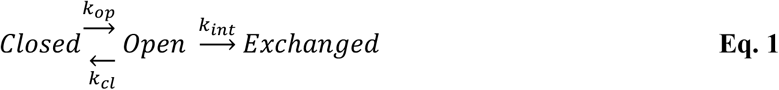

The *Closed* state itself makes not contribution to exchange. The reaction scheme can be related to the observed exchange rate *k_EX_* by [20]

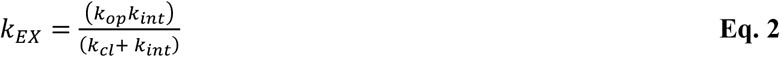

The equation has two limiting cases. In the EX2 (biomolecular exchange) limit at low pH, k_cl_ >> k_int_

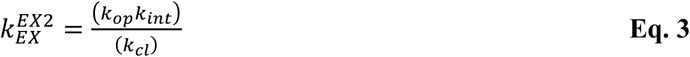

Since *k_int_* can be calculated using programs such as SPHERE [23], the observed exchange rate can be used to calculate [8, 9, 15, 20–22] the equilibrium constant for unfolding describing the concentrations of *Open* and *Closed* states *K_unf_* = (*k_op_*)/(*k_cl_*).

In the second limiting case EX1 (monomolecular exchange) at high pH, k_cl_ << k_int_ and Eq. 2 reduces to

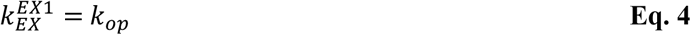

Thus, in the EX1 limit at high pH, the observable exchange rate depends on the opening rate describing the breaking of H-bonded structure to allow OH^-^-catalyzed exchange of exposed amide protons.

Simulations of the effects of different *k_cl_* rates calculated from Eq. 2 on *k_EX_*, with the *k_int_* rates obtained for Aβ(40) from the SPHERE program [23], are shown in Fig. 1B. The simulations show that the pH at which HX enters the EX1 limit, characterized by a pH-independent plateau, depends on the *k_cl_* rate. For a protein that folds without intermediates *k_cl_* can be equated to the protein folding rate since the entire structure is formed in a single highly cooperative reaction. Numerous kinetic protein folding studies have demonstrated that evolutionary constraints have selected for proteins to fold in a highly cooperative manner without intermediates and on extremely fast timescales [32–34]. A survey of folding rates for two-state proteins normalized to a uniform temperature of 25 °C, found values in a range between 0.5 x10^3^ and 2 x10^5^ s^-1^ [35]. A larger survey of 108 proteins similarly found an average protein folding rate of 0.4 x 10^3^ s^-1^ for two-state folders [36]. Thus, based on these folding rates, most typical proteins should be well in the EX1 exchange limit by about pH 10 to 11, according to the simulations in Fig. 1B.

Similar simulations (Supporting Information Figure S1) show that decreasing the unfolding rate *k_op_* while keeping *k_cl_* constant has the effect of shifting the pH-dependent exchange curves to slower *k_EX_* values. At low pH in the EX2 limit, decreasing *k_op_* while keeping *k_cl_* constant decreases K_op_ = k_op_/k_cl_ leading to slower exchange. At high pH in the EX1 limit, lower *k_op_* rates lead to slower *k_EX_* rates.

Thus, OH^-^-catalyzed exchange accelerates rates by a factor of 10 with every unit increase in pH. H-bonded structure protects amide protons even at high pH, however, leading to a plateau in the pH-dependence of exchange rates as the process becomes governed by H-bond breaking under EX1 conditions. Exchange protection due to H-bonded structure can be significant. For example, using H/D isotope exchange in D_2_O we measured *k_EX_* rates as slow as 10^-4^ s^-1^ in the proteins SN and LysN at pH 11.0 [21], a factor 10^9^-slower than the *k_int_* rates predicted from the amino acid sequences of the proteins. Even much lower protection allows amide protons participating in H-bonds to persist in NMR spectra obtained under alkaline conditions (pH > 10). Thus, high-pH NMR data should be useful for identifying amide protons participating in H-bonds.

### 2.2 Amide protons that survive at high pH accurately identify hydrogen bonds

As described above, protection from solvent exchange under EX1 conditions at high pH implies H-bonded structure. We reasoned that survival of amide protons in ^1^H-^15^N HSQC spectra at high pH might provide an alternative way to identify H-bonds in structures that are too unstable to give measurable solvent exchange protection in D_2_O, under EX2 conditions favored by neutral or slightly acidic pH. Compared to the more conventional H/D studies, to our knowledge there has not been an attempt to account for HX protection under EX1 conditions in terms of structural properties such as H-bounding. During the course of studies in our lab, we collected pH data on several proteins and obtained additional new data on others (Fig. 2, Figs. S2-S9, Tables S1-S10). This yielded a dataset of some 750 amide proton sites from 10 proteins with X-ray or cryoEM structures to investigate the relationship between HX protection at high pH and H-bonded structure. As illustrated in Fig. 2 for ubiquitin and the kinesin neck fragment comprised of an unfolded N-terminus and a C-terminal parallel coiled coil dimer, a subset of the ^1^H-^15^N HSQC correlations observed near neutral pH, survived in the spectrum under basic conditions at pH > 10.0. Similar results were observed for the three globular proteins CspA, LysN and SN, in an earlier publication [21]. Structural analyses showed that most of the protected amide protons were involved in protein H-bonding whereas unprotected amide protons did not participate in H-bonding. Exceptions in the data were amide protons not protected despite being involved in H-bonds, which we categorized as “False Negatives” (FN); and amide protons still detectable at high pH despite not being H-bonded, which we designated “False Positives” (FP). We expanded our NMR analysis to 10 proteins (Fig. 2, Fig. S2-S9) and analyzed the data on the persistence of amide protons at high pH in terms of H-bonding using X-ray, cryoEM, or NMR data for the proteins (Tables S1-S10).

**Figure 2.**
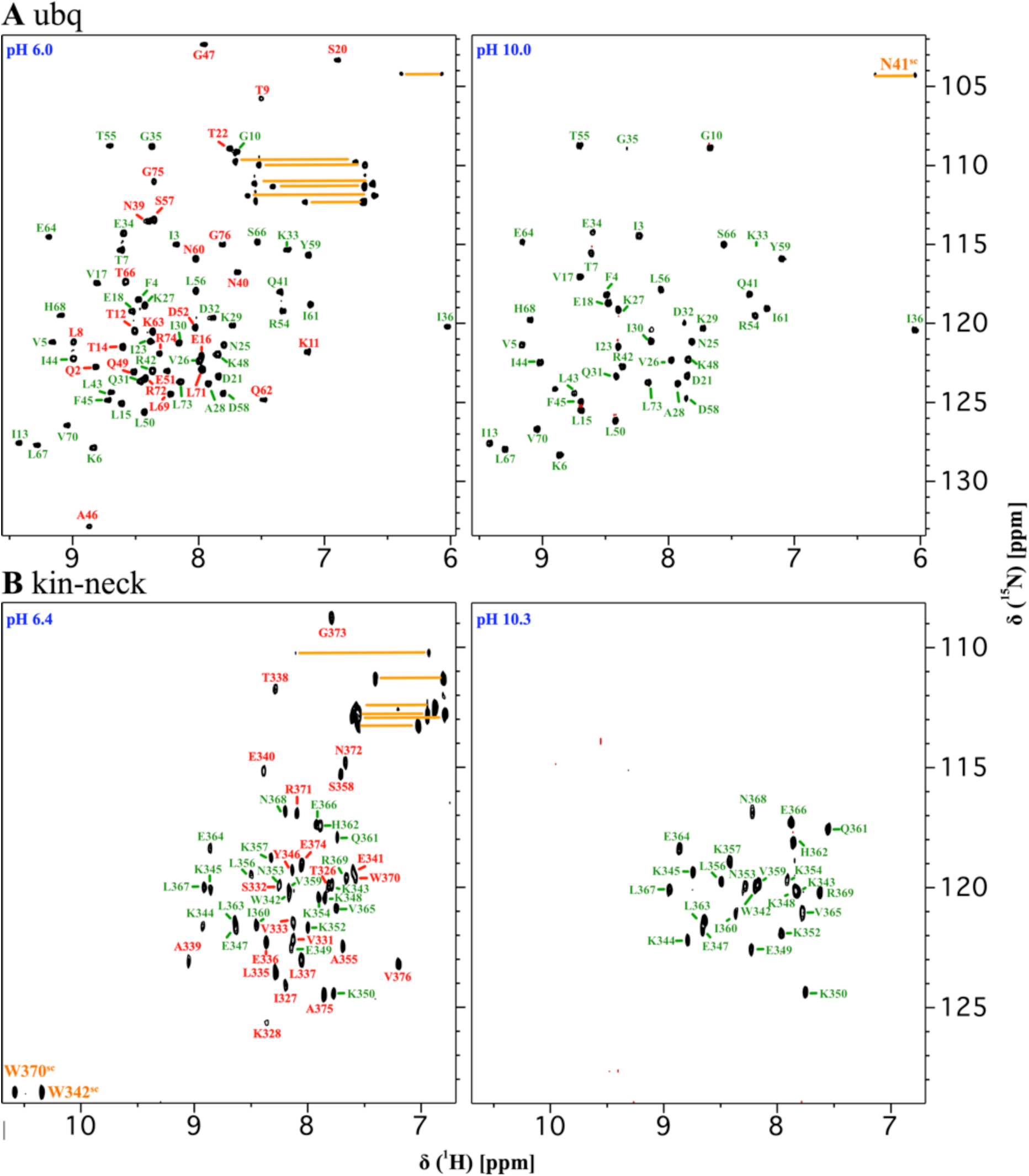
Survival of H-bonded amide protons at high pH. Representative data for **(A)** ubiquitin at pH 6.0 and 10.0, and (**B**) the kinesin neck fragment coiled coil at pH 6.4 and 10.3. The ubiquitin NMR spectra were obtained at a temperature of 25 °C while those for the kinesin neck coiled coil were at 37 °C. Peaks that survived at high pH are labeled green, those that did not are labeled red. NMR signals from sidechains are labeled orange. Additional data for 8 proteins are shown in Figs. S2-S9 and the relationship between amide proton NMR signal survival at high pH and H-bonding is summarized in Tables S1-S10.

The overall results for ∼750 amide protons in the 10 proteins are summarized in Table 1. The presence or absence of amide proton NMR signals between pH 10 and 11 can be used to ascertain H-bonds (protected amides) or the absence of H-bonds (unprotected amides) with ∼91% accuracy. About 6% of amide protons are FP, surviving at high pH even though they are not H-bonded. Some 3% are FN, not being detected at high-pH despite being H-bonded. Since all of the proteins studied also had data on isotope exchange protection in D_2_O at slightly acidic pH, we also performed the H-bonding analysis with this dataset (Table 1). The accuracy of H-bond identification from the traditional D_2_O protection data was 80%, slightly worse than that from the high-pH approach. The D_2_O data set had a higher proportion of both FP (8%) and FN (12%) amide protons. The fraction of FP amides may be due to the threshold to define D_2_O protection being set too low (typically survival of NMR signals after D_2_O incubation for 5 min-2 h, near 25 °C, at pH values between 5 and 6). Increasing the D_2_O incubation time would reduce cases of the FPs but increase the FNs as marginally stable structure would fall under the raised protection threshold. The higher proportion of FN amides in the D_2_O exchange data sets, is likely to reflect H-bonded structure that is too unstable to afford protection. This is exemplified by strand β5 of the protein CspA where protection cannot be observed in D_2_O, unless the protein is stabilized with 0.2 M concentrations of the osmolyte TMAO [15]. Indeed, the occurrence of such marginally stable structure was a primary factor motivating us to examine the use of amide proton survival in high-pH NMR as an alternative method for the identification of H-bonds.

**Table 1.**
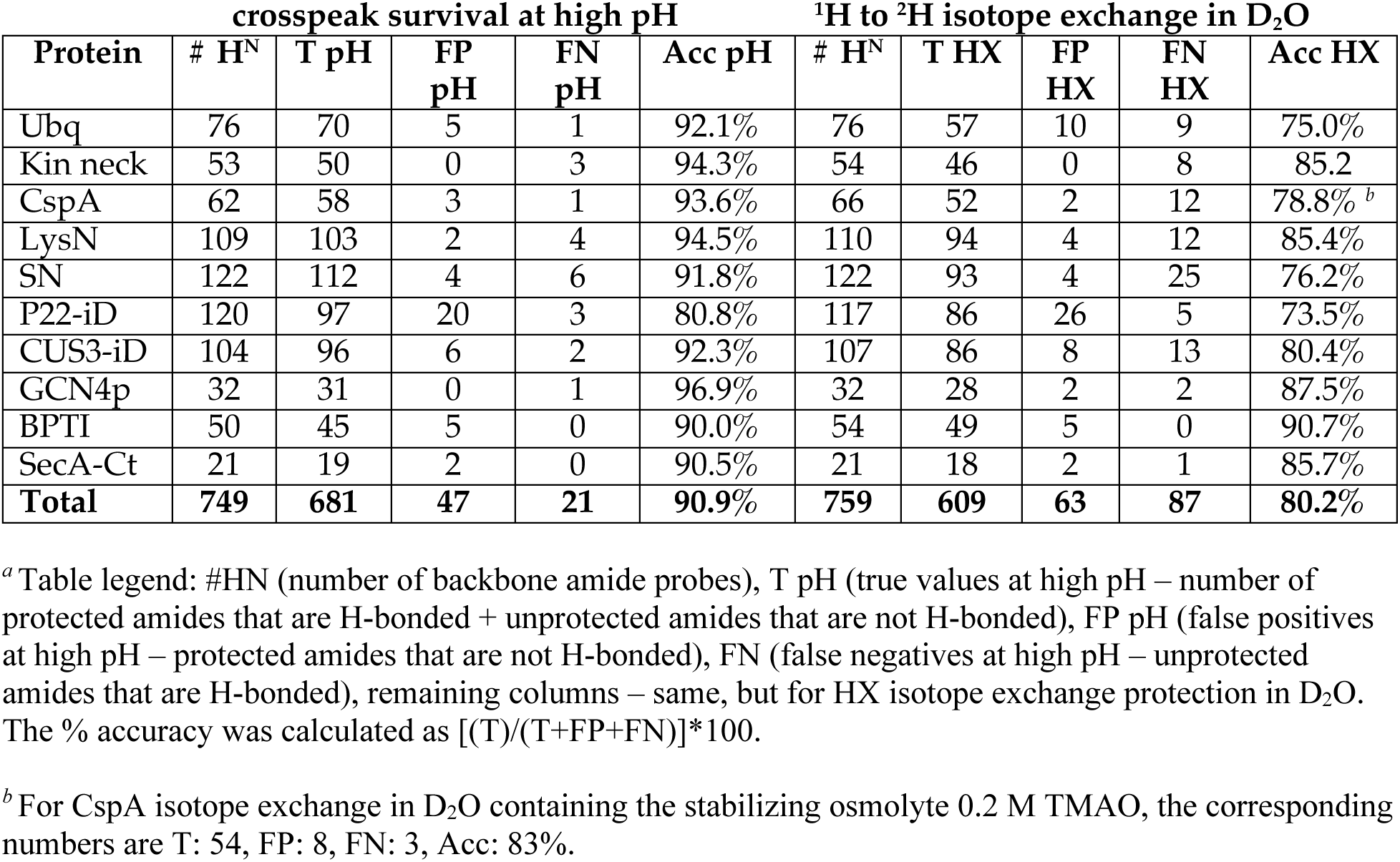
Accuracy of H-bond identification.

### 2.3 Foldon identification from amide proton survival in high-pH unfolding studies

In contrast to D_2_O exchange experiments under a single set of conditions, the high-pH approach is tunable across a range of exchange rates, from the slowest near neutral pH to the fastest at basic pH. This made us reason that looking at amide exchange protection over a series of increasing pH values would allow us to investigate the stability hierarchy of structural elements within a protein [37], up to the most stable secondary structure at alkaline pH before base-induced complete unfolding of the protein. As an alternative, we also explored the use of increasing temperature near pH 6 to identify the most persistent H-bonded structure. In addition to a higher likelihood of precipitation due to heating, thermal unfolding experiments had the drawback of giving a combination of ^1^H-^15^N HSQC NMR signals from the remaining structured molecules and unfolded molecules with amide protons that survived exchange at slightly acidic pH 5 to 6. In this regard, the high-pH ^1^H-^15^N HSQC experiments have the advantage of effectively filtering out signals from unstructured polypeptide segments through fast exchange. Above ∼pH 10, exchange rates are some thousand-fold larger than those compatible with detecting signals from unfolded proteins or segments of proteins (Fig. 1, at pH below 8), so only amide protons participating in stable H-bonded structure survive in the spectrum at pH 10-11.

Figure 3 shows representative ^1^H-^15^N HSQC pH titration data for two proteins: the GCN4p parallel coiled coil dimer [38], and the P22iD I-domain from the bacteriophage P22, that has a globular structure based on a six-stranded β-barrel with a smaller accessory subdomain [39]. As the sample pH is raised above pH 10, there is an increasing loss of amide protons from the spectra suggesting a progressive loss of structure. We infer that the amide protons surviving at the highest pH, correspond to the most persistent elements of H-bonded structure, and refer to these species as ‘foldons’ in analogy to partially unfolded quasi-independent units of H-bonded structure detected in native state hydrogen exchange (NSHX) experiments [9]. Whereas NSHX experiments detect foldons based on the stability of residual structure to unfolding near neutral pH under EX2 conditions [9], the foldons described in this work at high pH under EX1 conditions, correspond to the portions of the structure with the slowest unfolding rates. The distinction between folding stability and unfolding rates becomes effectively moot if protein folding rates are uniformly fast, as observed for several small proteins [35, 36], since then the unfolding rates would then be the principal determinant of folding stability. The interpretation of the disappearance of amide protons in terms of structural stability is supported amongst other data by comparison of the protein SN and its less stable fragment SN-OB. The SN-OB fragment retains the OB-fold structure of the parent protein SN but is missing the two C-terminal α-helices [40]. The structure and most of the sequence of the SN-OB fragment is the same as that of the conserved portion of the parent protein SN [40], but the stability of the SN-OB fragment is 2.4 Kcal/mol versus 6.4 Kcal/mol for SN [10]. Consistent with the differences in stability, at the same temperature of 25 °C, ^1^H-^15^N HSQC signals from amide protons in the SN-OB fragment can only be detected up to pH 10.2 while those from the more stable parent protein SN are observed up to pH 11.0 [21]. In addition to the aforementioned proteins GCN4p and P22iD, we obtained high-pH unfolding data for a second coiled coil dimer – the kinesin neck fragment, as well as the globular proteins CUS3id and ubiquitin. A summary of the pH values at which amide protons are last detected in the five proteins is given in Table S11. All five proteins show a progressive loss of amide protons with increasing pH, suggesting a stability hierarchy of partially unfolded states before complete unfolding at high pH. This is not the case for all of the proteins we looked at. For SN the surviving amides at pH 11.0 correspond to those H-bonded in the X-ray structure (Fig. S4, Table S5). By pH 11.3, all remaining amide protons are lost from the spectrum (not shown), in an apparently highly cooperative global unfolding process.

**Figure 3.**
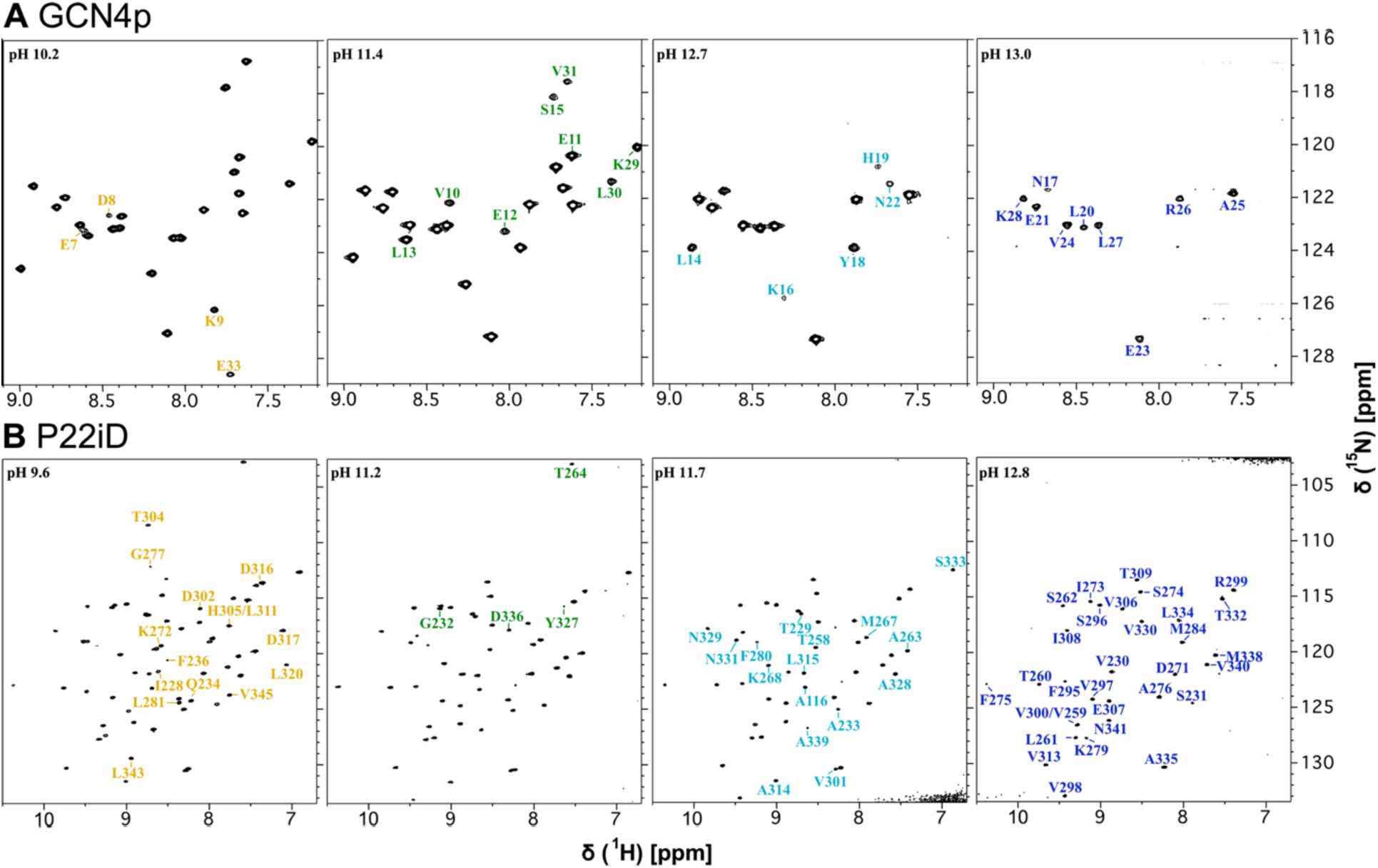
Selected high pH ^1^H-^15^N HSQC spectra used to identify residual H-bonded foldon structure. **(A)** Spectra of the GCN4p coiled coil from pH 10.2 to 13.0. (**B**) Spectra of the globular P22iD protein from pH 9.6 to 12.8. Each spectrum is annotated with labels for peaks that disappear above the stated pH. Above pH 13.0 for GCN4p and pH 12.8 for P22iD ^1^H-^15^N HSQC crosspeaks are no longer seen. The colors for the peak labels correspond to the pH ranges given in Fig. 4.

### 2.4 Foldons can consist of sequential or non-sequential structural motifs

To better understand the mechanisms of amide proton loss with increasing pH, we mapped the data (Table S11) on the structures of the 5 proteins studied (Fig. 4). For the GCN4p coiled coil there is a gradation from the N-terminus where amide protons survive only at moderately basic pH to the C-terminus where amid protons survive at the highest pH value (Fig. 4A). The 19-27 segment that survives up to pH 13.4 (the highest pH value before all NMR signals disappear), corresponds roughly to the ‘trigger site’ comprised of residues 18-31 [41]. Trigger sites are segments of coiled coil sequences that have the highest propensity for α-helical structure in the monomers, and are thought to thereby nucleate assembly and subsequent zippering of the coiled coil structure [41, 42]. It is interesting to note in this regard, that the segment of the protein that is the first to fold is also the last to unfold – both can be accounted for by the proclivity of the trigger site to form H-bonded α-helical structure.

**Figure 4.**
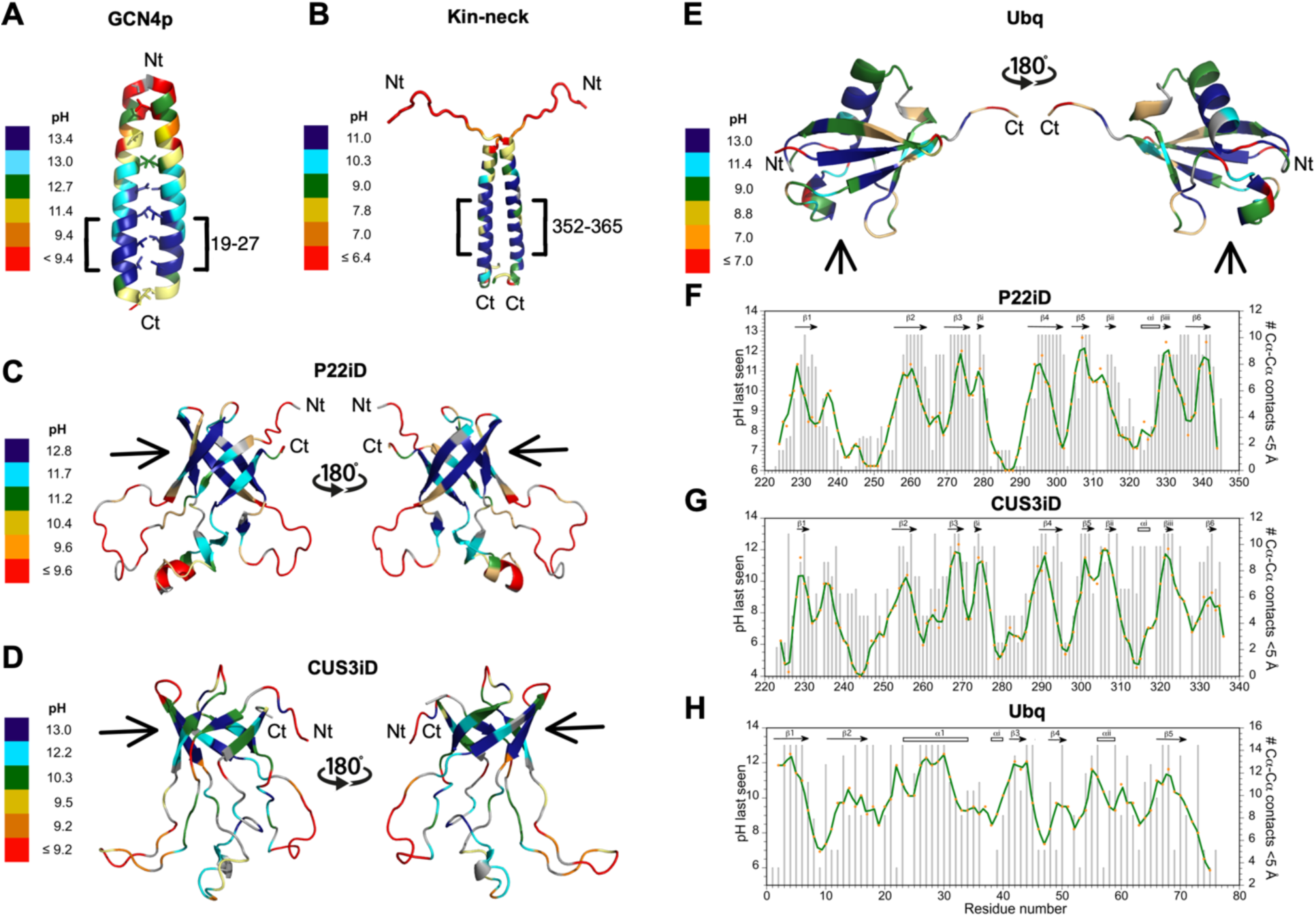
Structural analysis of amide proton persistence at high pH. Structural mapping of the pH values at which amide protons are last detected: **(A)** GCN4p coiled coil (PDB 2ZTA), (**B**) kinesin neck coiled coil fragment (PDB 8TT7), (**C**) P22iD (PDB 2M5S), (**E**) CUS3iD (PDB 6MNT), (**F**) ubiquitin (PDB 1UBQ). We used NMR structures of the isolated domains of P22iD and CUS3iD because the cryoEM structures of the corresponding domains form intact phage capsids have additional H-bonded structure involved in capsomer assembly. Note that the pH scales are different in each case since the proteins have distinct stabilities and unfolding cooperativities. Amides persisting at the highest pH values are indicated by brackets for the coiled coils (A,B) and arrows for the globular protein (C-E). Correlations of the pH values at which amide proton NMR signals are last observed (gray bars) with contact densities (green lines) for the globular proteins P22iD (**F**), CUS3iD (**G**) and ubiquitin (**H**). The contact densities (# of Cα-Cα contacts < 5 Å per residue) were smoothed by averaging over a 3-residue window.

For the second kinesin neck fragment coiled coil, there is also a gradation of amide proton NMR signal persistence from the rapidly exchanging N-terminus to the more exchange-resistant C-terminus (Fig. 4B). Similar to GCN4p, the 352-365 segment of the kinesin neck fragment with the amide protons that survive at the highest pH value of 11.0, score the highest for coiled coil preference with the DEEPCOIL1 program (https://toolkit.tuebingen.mpg.de/tools/deepcoil accessed on Jan 26, 2026; [43]). This region may correspond to the trigger site of the kinesin neck fragment.

For the globular protein P22iD, the most persistent amide protons map to non-contiguous segments comprising the six β-strands of a conserved antiparallel β-barrel structure (indicated by arrows in Fig. 4C). This foldon is similar to one identified in previous NSHX experiments of P22iD at pH 6.0 [22]. In the previous study, the six-stranded β-barrel had the greatest stability to unfolding while a subdomain consisting of a short α-helix and three short β-strands had a lower stability, consistent with the hierarchy of structure suggested by the high pH unfolding experiments in this work.

To test whether the pH dependence of amide protons is conserved in structurally related proteins we looked at the CUS3iD domain that has the same fold and function as P22iD [44] but only 35% sequence identity [45]. Like in P22iD, the six-stranded antiparallel β-barrel structure in CUS3iD (indicated by arrows in Fig. 4D) has the highest proportion of amide protons that survive at high pH, with the accessory subdomain consisting of three short β-strands and a one-turn α-helix, immediately below the β-barrel structure showing less persistence. The foldon structure consisting of the core of the six-stranded β-barrel, thus appears to be conserved between the structurally related P22iD and CUS3iD domains. Within the six-stranded β-barrel, strands 1 and 6 have a higher proportion of amide protons that exchange at lower pH in both P22iD and Cus3iD, suggesting a less firm connection between these strands. NMR studies support a disruption of the β1-β6 pairing of CUS3iD under acid-denaturing conditions that eventually leads to amyloid formation from the residual β-sheet structure of the partially open β-barrel (ATA, unpublished observations). We have not yet investigated if the β1-β6 connection that closes the β-barrel is also susceptible to disruption in P22iD, like in CUS3iD.

Finally, we collected high pH unfolding data for the well-studied protein ubiquitin (Fig. 4E). A number of partially folded states of ubiquitin have been reported including the A-state stabilized in alcohol under acidic conditions that only has the β1-β2 hairpin and long α-helix components of the structure [46], the related acidic U’ state with only the β1-β2 hairpin [47], the weakly populated acidic U” state that preserves most of the native structure but experiences an unfolding of residues 65-68 at the beginning of the C-terminal strand β5 [47], and a recently described pressure-denatured folding intermediate that involves an H-bond register shift in the β-sheet pairing of residues 65-73 from C-terminal strand β5 with strands β1 and β3 [48]. The high pH data for ubiquitin in this work does not support loss of any secondary structure elements, since all of those in the native structure have amides that survive at the highest pH: β1(residues 3-6), β2 (15-18), α (26-30), β3 (44-45), β4 (50), β5 (67-68). Together these persistent amides appear to form a ladder of secondary structure rungs through the portion of the globular structure closest to the N-terminus (arrow in Fig. 4E). By contrast the portion of the globular structure closer to the C-terminus has no amide protons that survive at high pH. This part of the molecule has a high fraction of basic Arg residues (not shown). Neutralization of the Lys residues between pH 10 and 11 may leave an excess of positively charged Arg residues at pH values > 12 that could destabilize structure in this region in a manner specific to ubiquitin at high pH, although this proposed mechanism will need further investigation.

Thus, denaturation experiments monitored by amide proton persistence at high pH show a variety of behaviors. For the two coiled coils GCN4p and kinesin neck, we see a gradual breakdown of structure within a single α-helical secondary structure. For the two related globular proteins P22iD and Cus3iD, the most persistent structure corresponds to the six β-strands that are not contiguous in the sequence but come together to form a conserved anti-parallel β-barrel structural motif. For ubiquitin, the most persistent structure maps to one side of the β-grasp folding motif that has a relatively neutral to negatively-charged electrostatic surface. Finally, for some proteins like SN, all H-bonded amides are lost over a narrow pH range, indicative of a cooperative global unfolding transition.

### 2.5 Density of inter-residue contacts is a determinant of foldon stability

In previous papers, we noted a correlation between hydrogen exchange stability in NSHX experiments and the density of inter-residue contacts in protein structures [21, 22]. This correlation extends even to amyloids probed by quenched HX, where the most protected amides in amyloids correspond to residues that form the largest number of contacts in the amyloid structure [49]. We see the same trend for the high-pH persistence of amide protons in the globular proteins studied in this work (Fig. 4 F-H). The R-values for correlations between the pH value at which amide protons are last observed and the number of Cα-Cα distance contacts in the structure < 5 Å, were 0.68 for P22iD, 0.66 for CUS3iD, and somewhat lower at 0.45 for ubiquitin. For all three proteins, the probability that these correlations could arise by chance is p <0.0001 at a 95% confidence level. Clearly, Cα-Cα contact density cannot be the only determining factor in the pH persistence of amide protons, since in the coiled coil examples the contact density should be nearly uniform through the structure (Fig. 4 A,B). As previously noted, the propensity of the polypeptide to fold into α-helix structure appears to be the dominant determinant in the high-pH HX behavior of coiled coils.

## 3. DISCUSSION

In this work we looked at the pH-dependence of NMR signals of ∼750 backbone amide protons from a database of 10 proteins. We found that the survival of amide proton NMR signals under basic conditions between pH 10 to 11 that promote fast HX can be used to identify amide proton H-bond donors with an accuracy of about 91 %. This compares favorably with the traditional method of inferring H-bonding from isotope exchange protection in D_2_O, which gave an accuracy of about 80% for the same dataset. The lower accuracy of the D_2_O isotope exchange method is due to a higher number of false negatives – amide protons that are H-bonded in marginally stable structure that offers insufficient protection to be detected in D_2_O. The alternative high-pH HX method has the advantage of being ‘tunable’ over a range of pH values, as long as these do not disturb the structure of the protein. Conversely, the advantage of the D_2_O exchange method is that the protein is studied under a fixed set of conditions where the only experimental variable is the substitution of D_2_O solvent for H_2_O.

Another method to identify protein H-bonds is exchange protection of amide protons at high temperature rather than high pH, however, this requires that the protein remains stable and won’t aggregate with temperature. Moreover, amide protons from unfolded sites can still persist at high temperatures, whereas exchange is accelerated by an order of magnitude with every unit increase in pH so that amides in unstructured regions are unlikely to make any contributions above about ∼pH 8.

Magnetization transfer from water to amide protons through exchange during the NOESY mixing time is another way to infer H-bonding status. We used this approach to distinguish a stably folded coiled coil from an unstructured IDP region in the protein IF1 [50]. The NOESY method, however, can only be used if exchange lifetimes are comparable to the ∼100 ms NOESY mixing time, and could be confounded by NOEs arising from the proximity of amide protons to protein-bound water molecules.

Finally, temperature coefficients, measured as the change in chemical shift of a nucleus with temperature (e.g. Δδ(H^N^)/ΔT) have been used to identify H-bonds both in folded proteins and loosely structured folding intermediates. In particular, Δδ(H^N^)/ΔT values more positive than -4.6 ppb/K predict H-bonds with an accuracy of 85-90% [22]. The accuracy can be further improved to as high as 95% by combining temperature coefficient data for multiple nuclei such as ^1^HN and ^15^N [22].

In our analysis of the relationship between amide proton protection at high-pH and H-bonding we observed examples of both false positives and false negatives. The false negatives occur because the stability of H-bonded structure can be insufficient to afford protection. This prompted us to explore the possibility of assessing the stability hierarchy of structure by shifting conditions to increasingly basic conditions where only the most persistent H-bonded structure survives. We used this approach for five proteins, and except for ubiquitin the residual structure observed to survive at the highest pH values generally agrees with that found in folding intermediates at neutral pH observed in NSHX experiments or protein fragment studies. This argues that the partially folded species observed at extremes of basic pH result from native-like interactions rather than representing an artefact of the specific alkaline denaturation conditions. The five proteins show differences in the pH values at which amide protons start to disappear as would be expected from the different stabilities of the structures involved, and in the range of pH values leading to a complete loss of signals due to the different unfolding cooperativities of the proteins.

False positives manifest exchange protection without apparent H-bonding, based on high-resolution structures of the respective proteins. It is generally accepted that participation in H-bonding rather than seclusion from solvent is the predominant requirement for amide proton exchange protection [26, 51]. Indeed, it is exceedingly rare for polar amide protons to occur in the interior of protein structures without being H-bonded [26]. Interestingly, examples of false positives in our high pH experiments often also correspond to false positives in the D_2_O exchange experiments as well. Examples include the amide protons of I36 and D58 for ubiquitin, as well as other amide protons in the nine other proteins we studied (Tables S1-S10). Possible explanations for the absence of H-bonding for these protected amides are errors in the X-ray or cryoEM structures, or contributions from H-bonding to solvent that are not discernable in the structures. Alternatively, studies on model compounds and proteins show that in a small number of cases amide protons lacking H-bonds can be protected by burial in the structure [52]. In our data, the fraction of false positive cases accounts for about 6-8% of the total (Table 1). This may be an overestimate, since some examples may be due to insufficiently stringent protection thresholds (pH too low or D_2_O incubation time too short) or the aforementioned structure errors in H-bonds detection.

For H-bonded sites protected in D_2_O exchange experiments, the degree of protection does not correlate with H-bonding strength as defined by H-bond distances, Δδ(H^N^)/ΔT temperature coefficients, or the magnitudes of NMR though-H-bond J-couplings [22, 53, 54]. Rather, the degree of protection appears to be correlated with the stability of the structure to unfolding as parametrized by the number of distance contacts with other residues in the structure [15, 21, 22, 55]. We found similar correlations in this work between the pH persistence of amide protons and the Cα-Cα contact density in the structure, at least for proteins with predominantly β-sheet structure (Fig. 4 F-H). For proteins with a coiled coil α-helical structure where the Cα-Cα contact density is uniform throughout the structure, the amide protons that persist at the highest pH values are those from the residues with the highest sequence propensity to adopt α-helical secondary structure.

In summary, we described methods to probe H-bonding and the unfolding dynamics of H-bonded structure using high-pH NMR. The approach does not require specialized labeling schemes or specialized NMR equipment and uses sensitive NMR experiments that can be done with unlabeled (2D ^1^H-TOCSY) or ^15^N-labled (2D ^1^H-^15^N HSQC) samples. A fingerprint ^1^H-^15^N HSQC spectrum for a protein at each pH value can be collected in a matter of minutes. Sensitivity can be further improved using ^1^H-^15^N SOFAST-HMQC, but since this experiment uses magnetization transfer through cross-polarization it will detect amide protons exchanging on faster timescales than those detected with INEPT transfer in HSQC spectra. A series of spectra at progressively increasing pH values can provide information on the H-bonded structure most resistant to alkaline unfolding. Since alkaline unfolding of the structure typically occurs at pH values that promote fast exchange of amide protons, unstructured polypeptides do not contribute to the spectrum. In the five proteins we examined in this work (except for perhaps ubiquitin), the residual structure at high pH is consistent with that observed in studies of partial unfolding near neutral pH. This indicates that unfolding at high pH occurs through non-specific structure disruption. The resulting foldon structures reflect the stability hierarchy of substructures, rather than partial unfolding specific to the conditions used to destabilize the protein. The resulting information on partially unfolded species observed in the high-pH unfolding studies could be relevant to understanding folding mechanisms, protein misfolding related to disease, the stability of pharmaceutical proteins, and conformational transitions important in protein function.

## 4. MATERIALS AND METHODS

### 4.1 Protein samples

Unless otherwise noted all the NMR spectra in this study were obtained at a temperature of 25 °C, with protein concentrations ranging from 0.1 to 1 mM. Exceptions were studies of kin-neck at 37 °C, and BPTI at 30 °C. Protein samples for NMR were prepared by *E coli* expression and purified according to previously published conditions as follows: Kin-neck [56], CspA [15, 57], LysN [58–60], SN [21], GCN4p [38], P22iD [39], Cus3iD [45]. Ubiquitin was expressed in *E. coli* BL21(DE3) cells using a pET28b(+) vector from Novagen (Madison, WI), with the ubiquitin gene inserted via the *Nhe*1 and *Not*I restriction sites. Uniform ^15^N-isotope labeling of ubiquitin and its purification were done as previously described [61]. ^15^N-labeled Aβ(40) was from rPeptide (Watkinsville, GA). BPTI was purchased as bovine lung aprotinin (Cat Number 616370-M, ≥ 95% pure) from Sigma (St. Louis, MO). SecA-Ct was custom synthesized to ≥ 98% purity by Biopeptide (San Diego, CA) and studied in the presence of 1:1 ZnCl_2_ under preciously described sample conditions [62]. Sample pH values were adjusted with small (0.3-3 µL) aliquots of 1N HCl and NaOH. A Mettler (Columbus OH) MA235 pH meter equipped with a glass InLab pH electrode was used for all pH measurement. The sample pH was taken as the average of measurements before and after NMR experiments, which typically differed by < 0.1 pH unit.

### 4.2 NMR data acquisition and analysis

NMR assignments at the various pH values employed in this study were extended with 2D titration experiments using ^1^H-^15^N HSQC (pulse sequences *gNhsqc* for Varian and *fhsqcf3gpph* for Bruker) or TOCSY (Bruker pulse sequence *dipsi2gpph19*) spectra, starting from published NMR assignments near neutral pH. Chemical shift assignment files and annotated fingerprint regions of NMR spectra were obtained from the BioMagResBank (BMRB) and publications as follows: ubq (BMRB 6457, [63]), kin-neck (BMRB 52075, [56]), CspA (BMRB 4296, [64]), LysN ([58]), SN (BMRB 4052, [65]), GCN4p (BMRB 30027, [38]), P22iD (BMRB 18566, [39]), CUS3iD (BMRB 25263, [45]), BPTI (BMRB 4968), SecA-Ct (BMRB 6289). Data on protection from ^1^H/^2^H isotope exchange in D_2_O was obtained from the literature under the following conditions: ubq (1 h, pH 3.5, 22 °C, [66]), kin-neck (0.5 h, pH 6.1, 37 °C, [56]), CspA (3 min, pH 6.0, 30 °C, [64]), LysN (40 min, pH 6.0, 20 °C, [59]), SN (1 h, pH 5.2, 25 °C, [21]), GCN4p (3 min, pH 7.0, 6 °C, [67]), P22iD (2 h, pH 6.0, 37 °C, [39]), CUS3iD (2 h, pH 6.0, 25 °C, [45]), BPTI (100 min, pH 3.5, 36 °C, [68]), SecA-Ct (5 min, pH 6.8, 25 °C, [62]). H-bonds were calculated with the program HBPLUS [69] from structures with the following PDB codes: ubq (1UBQ), kin-neck (3KIN), CspA (1MJC), LysN (1BBU_A), SN (1STN), GCN4p (2ZTA_A), P22iD (8I1V_A and [53]), CUS3iD (8SKG_A and [53]), BPTI (1G6X), SecA-Ct (1TM6). Calculations of distance contacts between residues were done with the MAPIYA server using a contact cutoff of 5 Å and an intra-contact filter set to zero (https://mapiya.lcbio.pl accessed on Jan 18, 2026; [70]). Intrinsic HX rates were calculated from amino acid sequences with the program SPHERE (https://spin.niddk.nih.gov/bax/nmrserver/sphere/ accessed on Dec 21, 2025; [23]).

## AUTHOR CONTRIBUTIONS

**A.T. Alexandrescu:** Conceptualization; project administration, supervision; methodology, resources; investigation; formal analysis, visualization; writing – original draft, review and editing.

**A. J. Rua:** investigation; formal analysis, writing – review and editing

**S. Shah:** formal analysis, writing – review and editin

**D. Fairchild:** resources, writing – review and editing

**I. Bezsonova:** resources, writing – review and editing

## SUPPLEMENTARY DATA

Nine figures showing high pH NMR data for eight proteins, and simulations of the pH dependence of HX. Eleven tables showing relationship between H-bonding and survival of amide proton NMR signals at high pH for ten proteins.

## Abbreviations

Aβ(40): 40-residue amyloid beta peptide
H-bond: hydrogen bond
H/D: hydrogen/deuterium
HSQC: heteronuclear single quantum correlation
HX: hydrogen exchange
NMR: nuclear magnetic resonance
NSHX: native state hydrogen exchange
*k_cl_*: closing rate
*k_int_*: intrinsic exchange rate
*k_op_*: opening rate
*K_unf_*: equilibrium constant for protein unfolding.

