## Supplemental Material for "High-pH NMR to Identify Macromolecular Hydrogen-Bonds and Foldons"

**Table of contents:**

**Figure S1 – HX simulations at constant closing ($k_{cl}$) and decreasing opening ($k_{op}$) rates**
**Figure S2 – CspA: loss of $^1$H-$^{15}$N HSQC signals between pH 6.5 and 10.6**
**Figure S3 – LysN: loss of LysN $^1$H-$^{15}$N HSQC signals between pH 6.0 and 11.0**
**Figure S4 – SN: loss of $^1$H-$^{15}$N HSQC signals between pH 6.5 and 11.0**
**Figure S5 – GCN4p: loss of $^1$H-$^{15}$N HSQC signals between pH 6.2 and 9.9**
**Figure S6 – P22iD: loss of $^1$H-$^{15}$N HSQC signals between pH 6.2 and 10.4**
**Figure S7 – Cus3iD: loss of $^1$H-$^{15}$N HSQC signals between pH 6.1 and 10.3**
**Figure S8 – BPTI: loss of 2D $^1$H-TOCSY signals at 30 °C between pH 6.0 and 10.6**
**Figure S9 – SecA-Ct: loss of 2D $^1$H-TOCSY signals between pH 7.4 and 10.8**

**Table S1 Ubiquitin: H-bonding and surviving $^1$H-$^{15}$N HSQC correlations at pH 10.0**
**Table S2 Kin-neck: H-bonding and surviving $^1$H-$^{15}$N HSQC correlations at pH 10.3**
**Table S3 CspA: H-bonding and surviving $^1$H-$^{15}$N HSQC correlations at pH 10.6**
**Table S4 LysN: H-bonding and surviving $^1$H-$^{15}$N HSQC correlations at pH 11.0**
**Table S5 SN: H-bonding and surviving $^1$H-$^{15}$N HSQC correlations at pH 11.0**
**Table S6 GCN4p: H-bonding and surviving $^1$H-$^{15}$N HSQC correlations at pH 9.9**
**Table S7 P22iD: H-bonding and surviving $^1$H-$^{15}$N HSQC correlations at pH 10.4**
**Table S8 Cus3iD: H-bonding and surviving $^1$H-$^{15}$N HSQC correlations at pH 10.3**
**Table S9 BPTI: H-bonding and surviving HN-H$\alpha$ TOCSY correlations at pH 10.6**
**Table S10 SecA-Ct: H-bonding and surviving HN-H$\alpha$ TOCSY correlations at pH 10.8**
**Table S11 Sequential high pH unfolding of 5 proteins characterized by pH values above which amide peaks are no longer seen**

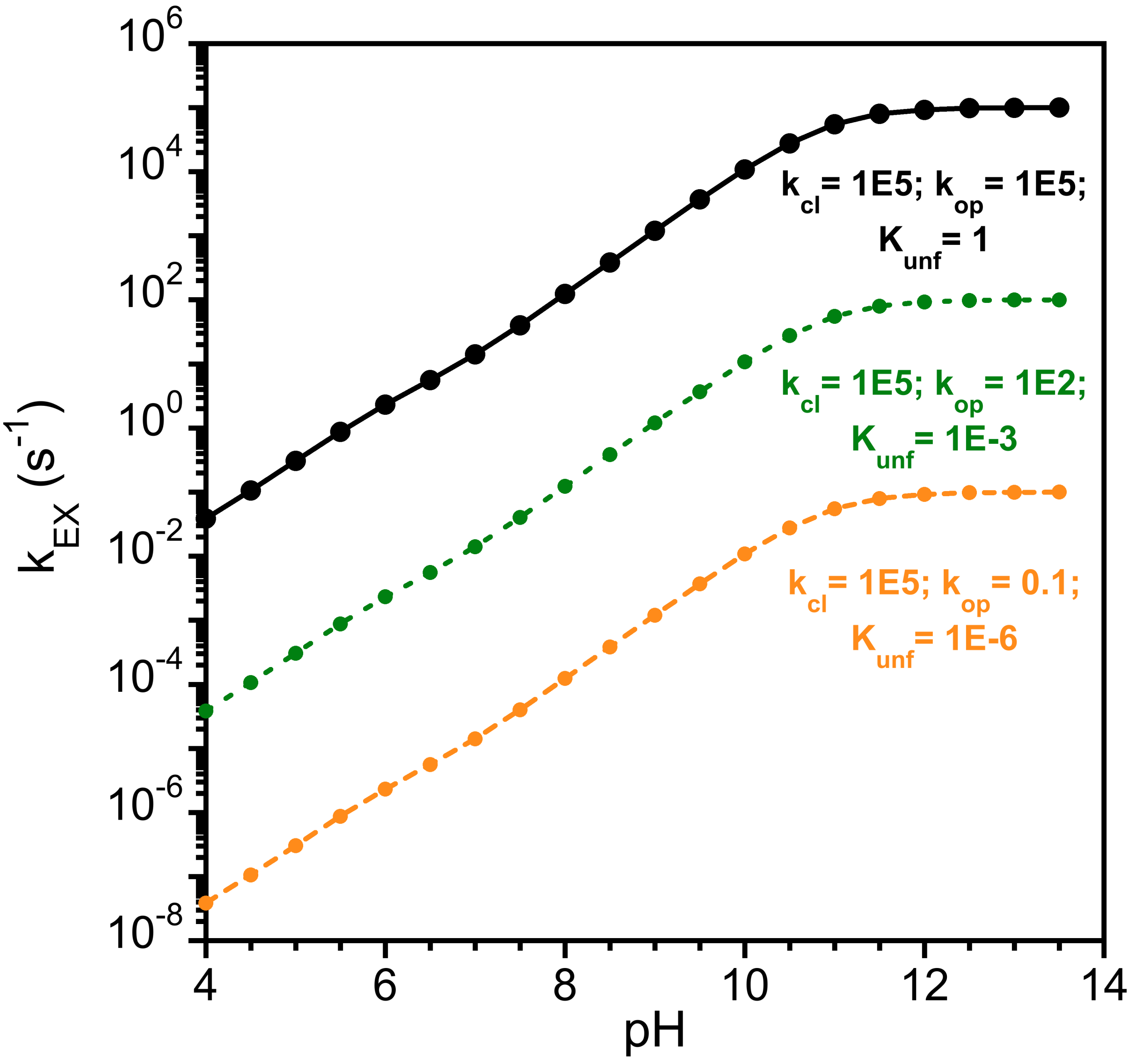

**Figure S1 – HX simulations at constant closing ($k_{cl}$) and decreasing opening ($k_{op}$) rates.** Starting with intrinsic rates calculated for the Aβ(1-40) sequence at 20 °C in $H_2O$ with the program SPHERE (Fig. 1), we calculated pH-dependent HX profiles using Eq. 1 and the parameters given in the figure. Note that in all three simulations the EX1 plateau occurs at pH 10.5, since the position of the plateau depends on the closing rate $k_{cl}$ which is fixed to $10^5$ $s^{-1}$. With the closing rate ($k_{cl}$) kept constant, the curves shift downwards (green, orange) to slower exchange rates as $k_{op}$ is decreased. At low pH approaching the EX2 limit, the equilibrium constant for unfolding $K_{unf}$ = ($k_{op}$/$k_{cl}$) decreases causing slower HX. At high pH values above 10.5 in the EX1 limit plateau, HX depends on the opening rate $k_{op}$ which is decreased in the simulations. If $k_{cl}$ were increased the curves would have similar shapes but the EX1 plateaus would shift to lower pH values (see Fig. 1).

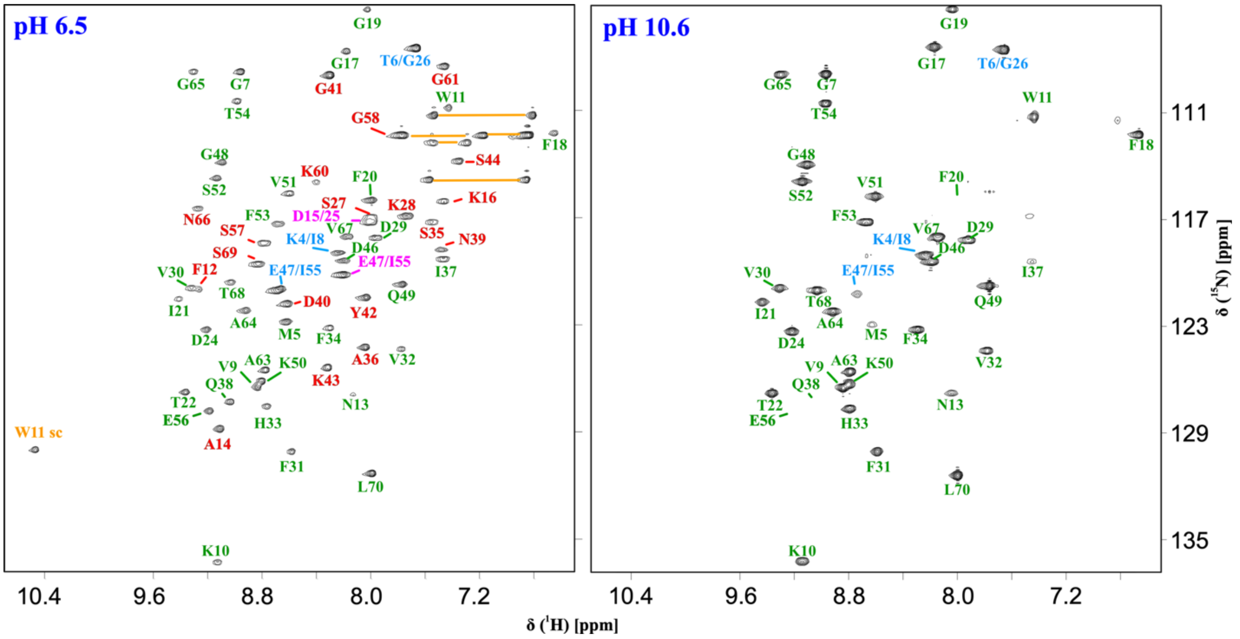

**Figure S2 – CspA: loss of $^{1}$H-$^{15}$N HSQC signals between pH 6.5 and 10.6.** For this figure and Fig. S3 to S9 the labels are color-coded as follows: green – amide NMR signals that survive at high pH, red – amide NMR signals not detected at high pH, light blue – overlapping NMR signals, orange – NMR signals from sidechains.

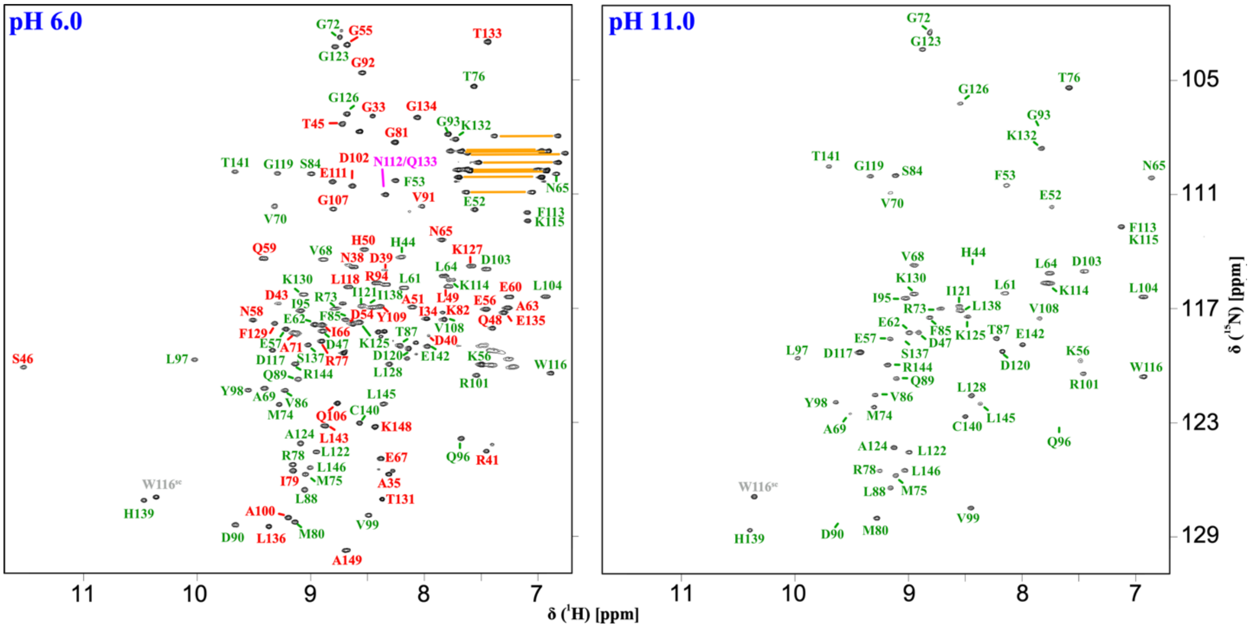

**Figure S3 – LysN: loss of $^{1}$H-$^{15}$N HSQC signals between pH 6.0 and 11.0.** Please see Fig. S2 for label color legend.

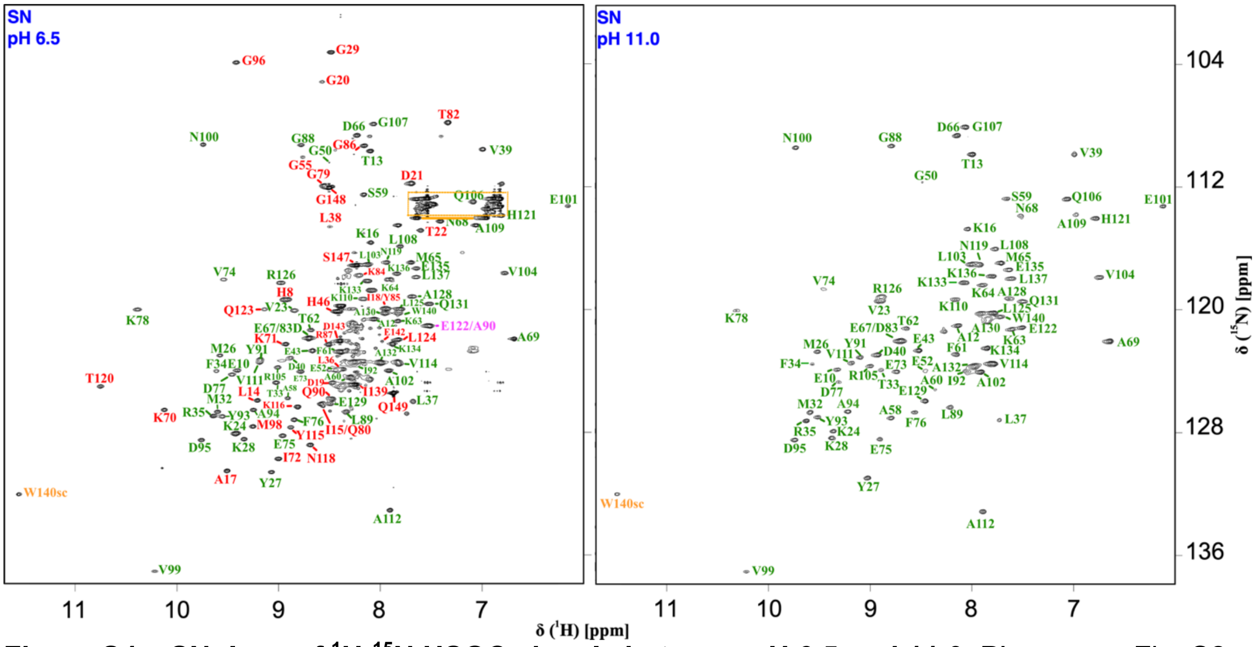

**Figure S4 – SN: loss of $^1$H-$^{15}$N HSQC signals between pH 6.5 and 11.0.** Please see Fig. S2 for label color legend.

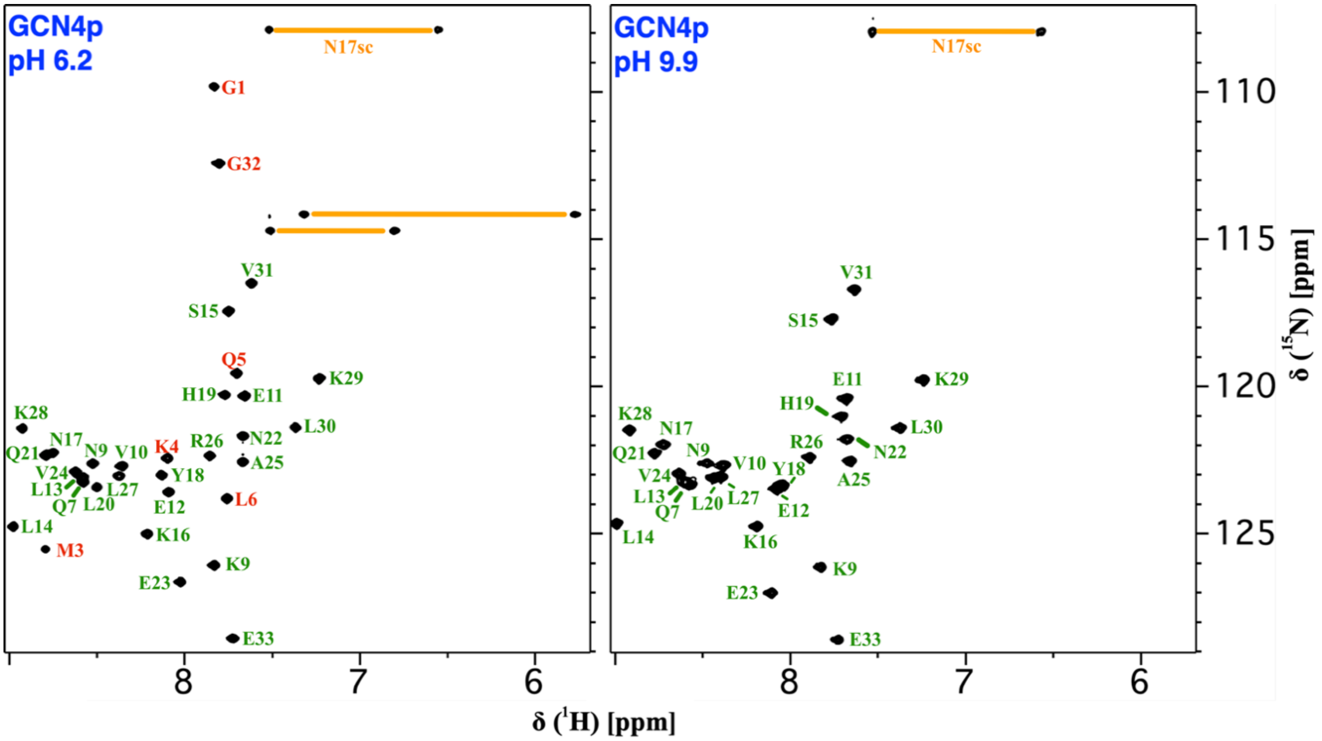

**Figure S5 – GCN4p: loss of $^1$H-$^{15}$N HSQC signals between pH 6.2 and 9.9.** Please see Fig. S2 for label color legend.

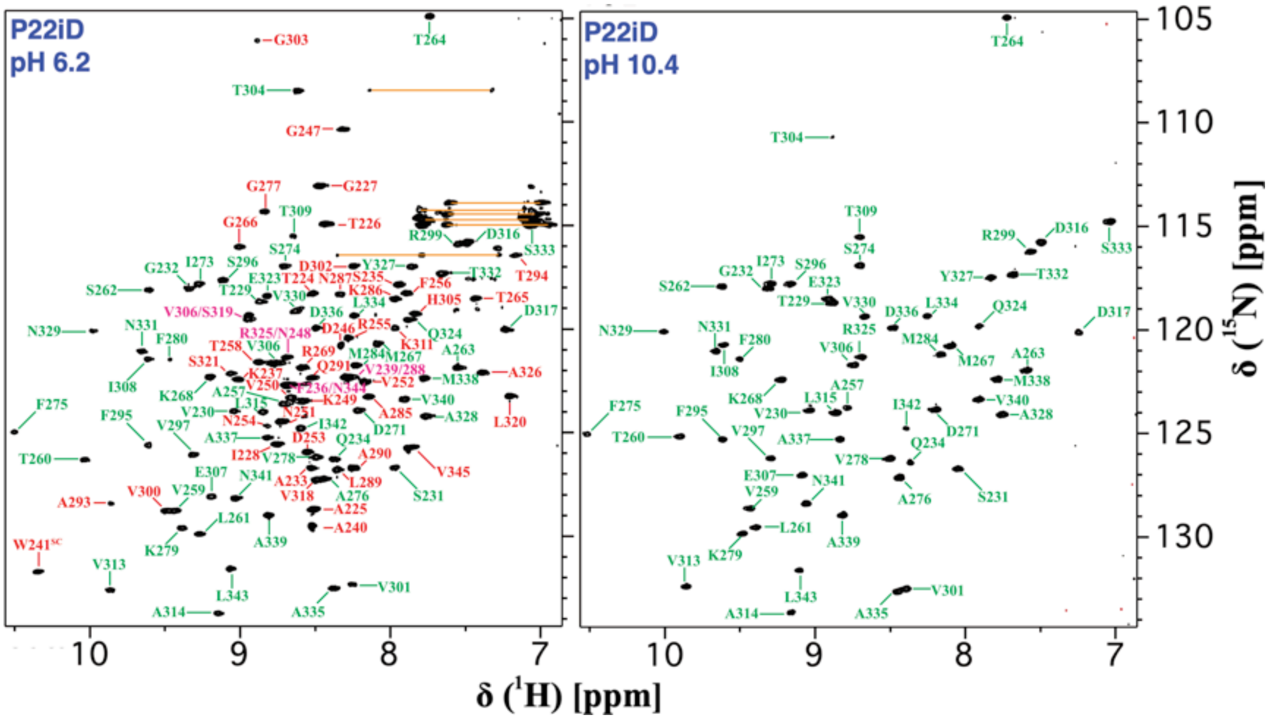

**Figure S6 – P22iD: loss of $^{1}H$-$^{15}N$ HSQC signals between pH 6.2 and 10.4.** Please see Fig. S2 for label color legend.

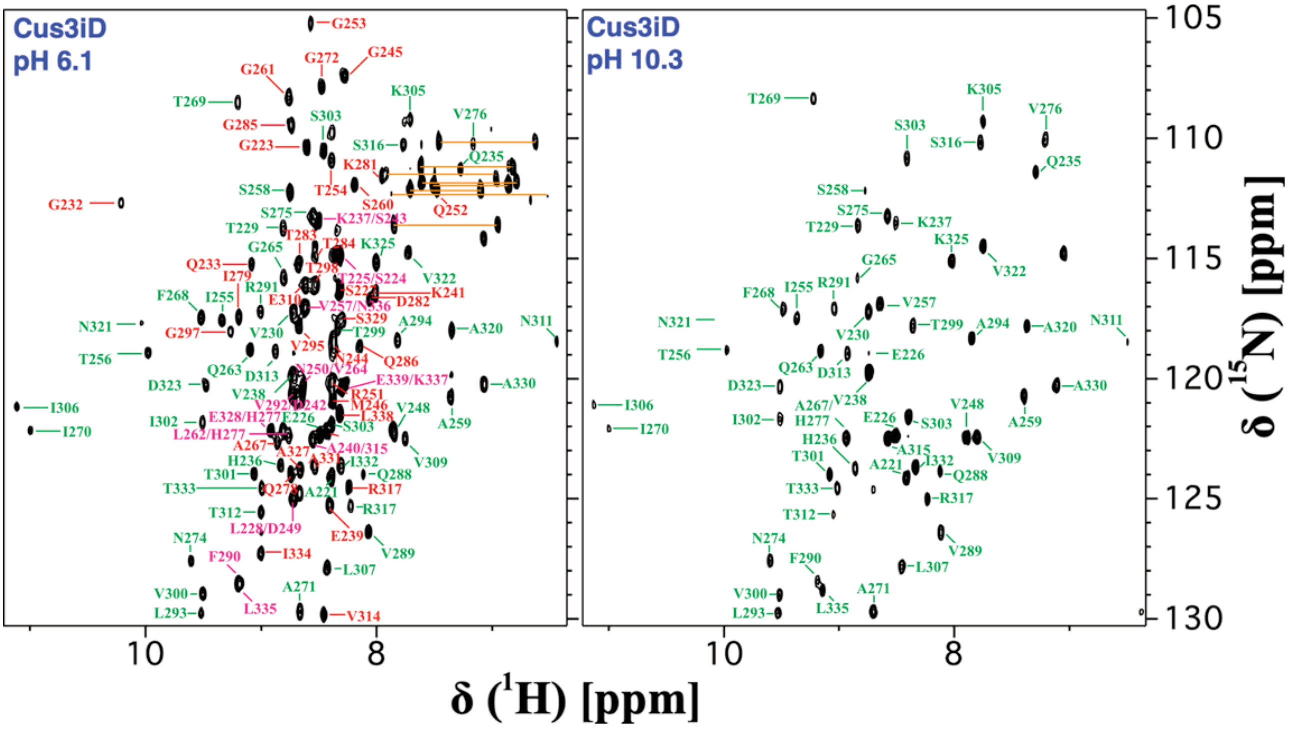

**Figure S7 – Cus3iD: loss of $^{1}H$-$^{15}N$ HSQC signals between pH 6.1 and 10.3.** Please see Fig. S2 for label color legend.

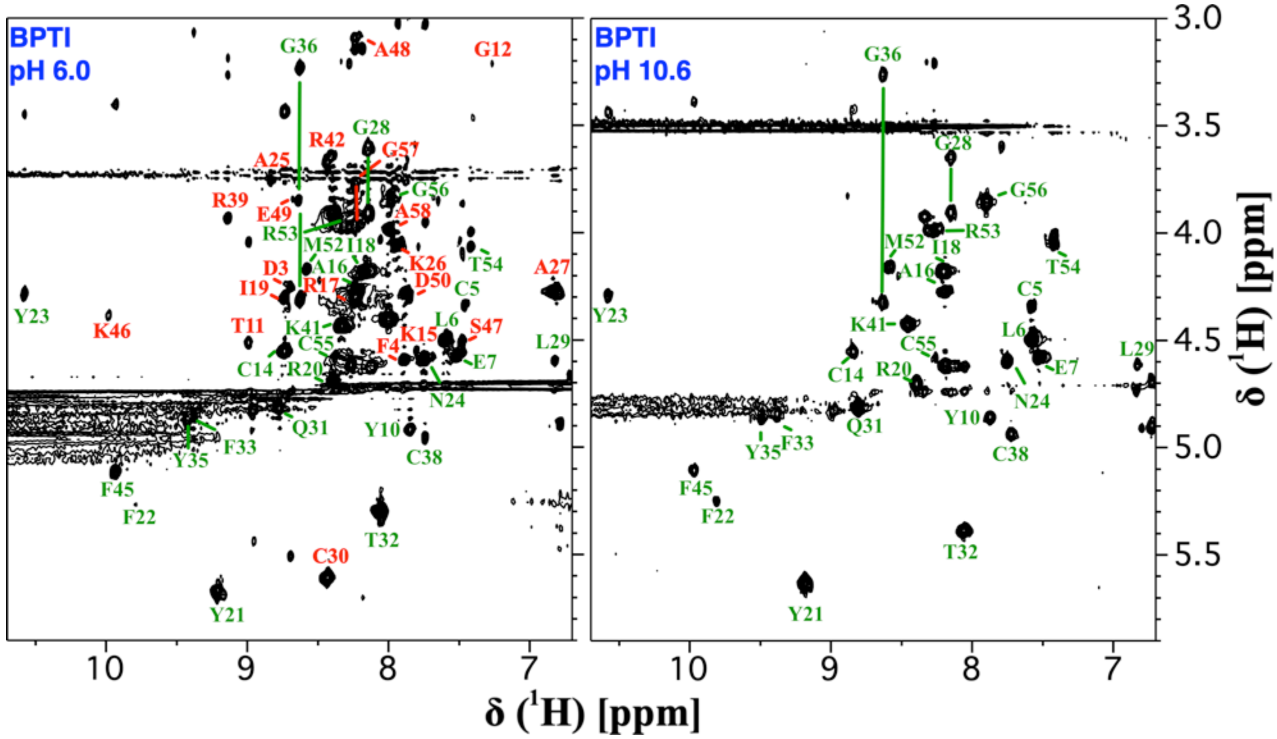

**Figure S8 –BPTI: loss of 2D [1]H-TOCSY signals at 30 °C between pH 6.0 and 10.6.** Please see Fig. S2 for label color legend.

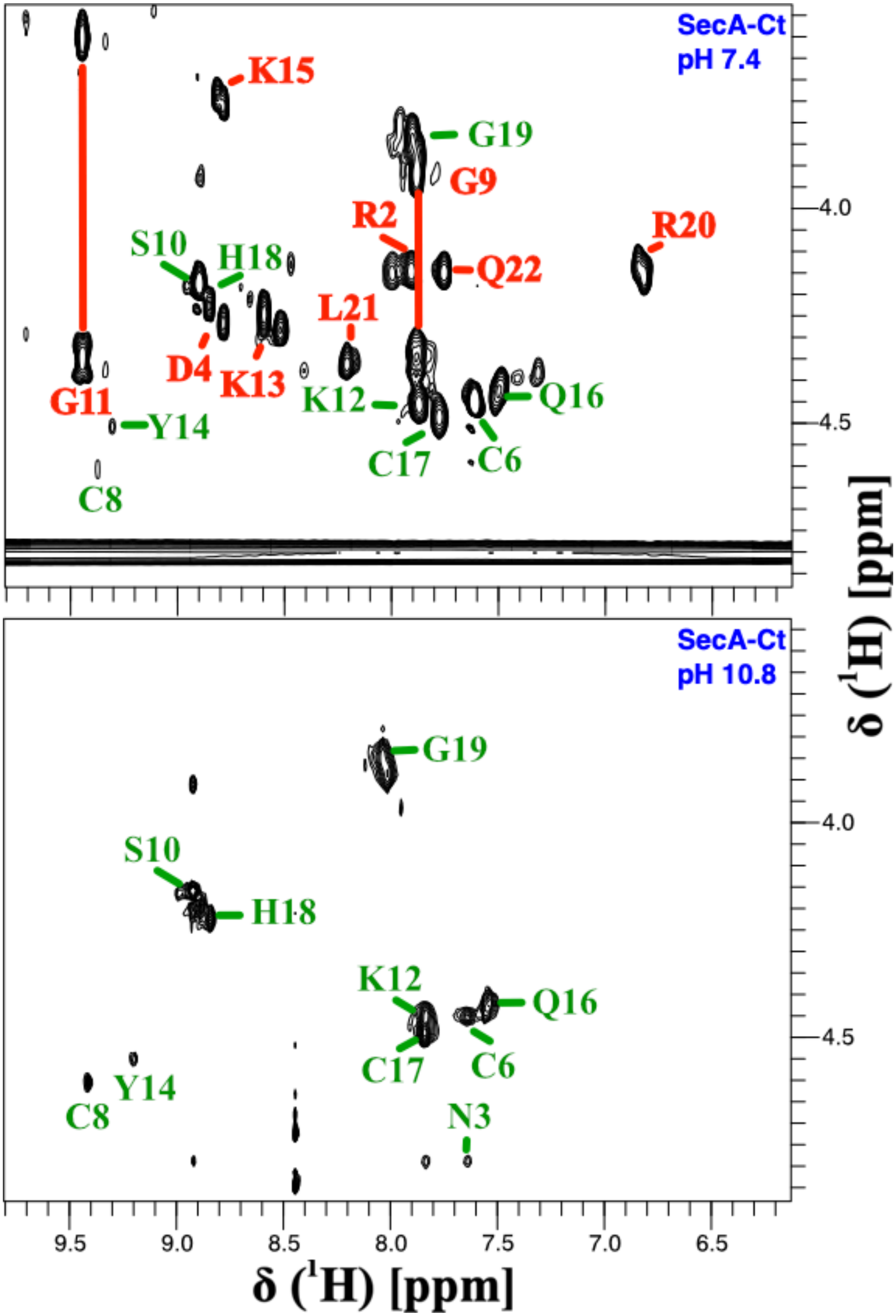

**Figure S9 – SecA-Ct: loss of 2D $^{1}$H-TOCSY signals between pH 7.4 and 10.8.** Please see Fig. S2 for label color legend.

**Table S1 Ubiquitin: H-bonding and surviving $^{1}H$-$^{15}N$ HSQC correlations at pH 10.0.**[a]

| # | aa | $H^N$ | N | notes | HB | hi-pH | HX | T/F hi-pH | T/F HX |
|---|---|---|---|---|---|---|---|---|---|
| 1 | M | | | | | n | n | T | T |
| 2 | Q | | | | | n | y | T | FP |
| 3 | I | 8.24 | 114.5 | | L15 | y | y | T | T |
| 4 | F | 8.50 | 118.2 | | S65 | y | y | T | T |
| 5 | V | 9.16 | 121.5 | | I13 | y | y | T | T |
| 6 | K | 8.87 | 128.4 | | L67 | y | y | T | T |
| 7 | T | 8.61 | 115.6 | | K11 | y | y | T | T |
| 8 | L | | | | | n | n | T | T |
| 9 | T | | | | | n | n | T | T |
| 10 | G | 7.67 | 109.0 | | T7 | y | n | T | FN |
| 11 | K | | | | | n | n | T | T |
| 12 | T | | | | | n | y | T | FP |
| 13 | I | 9.42 | 127.6 | | V5 | y | y | T | T |
| 14 | T | | | | | n | n | T | T |
| 15 | L | 8.69 | 125.5 | | I3 | y | y | T | T |
| 16 | E | | | | | n | y | T | FP |
| 17 | V | 8.71 | 117.1 | | M1 | y | y | T | T |
| 18 | E | 8.49 | 118.8 | | D21 OD2 | y | n | T | FN |
| 19 | P | | | | | n | n | T | T |
| 20 | S | | | | | n | n | T | T |
| 21 | D | 7.86 | 123.4 | | E18 | y | n | T | FN |
| 22 | T | | | | | n | y | T | FP |
| 23 | I | 8.40 | 121.6 | | R54 | y | y | T | T |
| 24 | E | | | | | n | n | T | T |
| 25 | N | 7.82 | 121.2 | | T22 OG1 | y | y | T | T |
| 26 | V | 7.99 | 122.4 | | T22 | y | y | T | T |
| 27 | K | 8.41 | 119.2 | | I23 | y | n | T | FN |
| 28 | A | 7.94 | 123.8 | | E24 | y | y | T | T |
| 29 | K | 7.73 | 120.3 | | N25 | y | y | T | T |
| 30 | I | 8.14 | 121.2 | | V26 | y | y | T | T |
| 31 | Q | 8.42 | 123.4 | | K27 | y | y | T | T |
| 32 | D | 7.88 | 120.0 | weak | A28 | y | y | T | T |
| 33 | K | 7.39 | 116.0 | weak | K29 | y | n | T | FN |
| 34 | E | 8.61 | 114.3 | | I30 | y | n | T | FN |
| 35 | G | 8.35 | 108.9 | weak | Q31 | y | n | T | FN |
| 36 | I | 6.06 | 120.4 | | | y | y | FP | FP |
| 37 | P | | | | | n | n | T | T |
| 38 | P | | | | | n | n | T | T |
| 39 | D | | | | | n | n | T | T |
| 40 | Q | | | | | n | y | T | FP |
| 41 | Q | 7.37 | 118.1 | | P38 | y | y | T | T |

| | | | | | | | | | |
|---|---|---|---|---|---|---|---|---|---|
| 42 | R | 8.36 | 122.8 | | V70 | y | y | T | T |
| 43 | L | 8.75 | 124.5 | | | y | n | FP | T |
| 44 | I | 9.01 | 122.5 | | H68 | y | y | T | T |
| 45 | F | 8.70 | 125.1 | | K48 | y | y | T | T |
| 46 | A | | | | | n | n | T | T |
| 47 | G | | | | | n | n | T | T |
| 48 | K | 7.85 | 122.3 | | F45 | y | y | T | T |
| 49 | Q | | | | | n | y | T | FP |
| 50 | L | 8.43 | 126.2 | | L43 | y | y | T | T |
| 51 | E | | | | | n | n | T | T |
| 52 | D | | | | | n | n | T | T |
| 53 | G | | | | | n | n | T | T |
| 54 | R | 7.32 | 119.6 | | E51 | y | n | T | FN |
| 55 | T | 8.72 | 108.8 | | D58 OD1 | y | y | T | T |
| 56 | L | 8.07 | 117.9 | | D21 | y | y | T | T |
| 57 | S | | | | P19 | n | y | FN | T |
| 58 | D | 7.86 | 124.8 | weak | | y | y | FP | FP |
| 59 | Y | 7.12 | 116.0 | | L56 | y | y | T | T |
| 60 | N | | | | | n | y | T | FP |
| 61 | I | 7.23 | 119.1 | | L56 | y | y | T | T |
| 62 | Q | | | | | n | n | T | T |
| 63 | K | | | | | n | n | T | T |
| 64 | E | 9.16 | 114.9 | | Q2 | y | n | T | FN |
| 65 | S | | | | Q62 | y | y | T | T |
| 66 | T | 7.56 | 115.1 | | | n | n | T | T |
| 67 | L | 9.30 | 128.0 | | F4 | y | y | T | T |
| 68 | H | 9.11 | 119.8 | | I44 | y | y | T | T |
| 69 | L | | | | K6 | y | y | T | T |
| 70 | V | 9.05 | 126.8 | | R42 | y | y | T | T |
| 71 | L | | | | | n | n | T | T |
| 72 | R | | | | | n | n | T | T |
| 73 | L | 8.17 | 123.8 | | | y | n | FP | T |
| 74 | R | | | | | n | n | T | T |
| 75 | G | | | | | n | n | T | T |
| 76 | G | | | | | n | n | T | T |

[a] Columns legend: # (residue number in the sequence); aa (amino acid type), HN (HN chemical shift of amide protons surviving at high pH), N (N chemical shift of amide protons surviving at high pH), HB (hydrogen bonds calculated from the PDB structures given in the Methods section), hi-pH (crosspeak survives at high pH), HX (crosspeak is protected in $D_2O$ near neutral pH under the conditions given in the Methods section), TF hi-pH (true if high pH data matches presence/absence of H-bonding, otherwise false positive or false negative), TX HX (true if HX protection near neutral pH matches presence/absence of H-bonding, otherwise false positive or false negative). FP (false positive) – peak is protected but not H-bonded, FN (false negative)– peak is not protected but H-bonded.

**Table S2 Kin-neck: H-bonding and surviving $^{1}H$-$^{15}N$ HSQC correlations at pH 10.3.**[a]

| # | aa | $H^N$ | N | notes | HB | hi-pH | HX[a] | T/F hi-pH | T/F HX |
|---|---|---|---|---|---|---|---|---|---|
| 323 | G | | | | | n | n | T | T |
| 324 | S | | | | | n | n | T | T |
| 325 | K | | | | | n | n | T | T |
| 326 | T | | | | | n | n | T | T |
| 327 | I | | | | | n | n | T | T |
| 328 | K | | | | | n | n | T | T |
| 329 | N | | | | | n | n | T | T |
| 330 | T | | | | | n | n | T | T |
| 331 | V | | | | | n | n | T | T |
| 332 | S | | | | | n | n | T | T |
| 333 | V | | | | | n | n | T | T |
| 334 | N | | | | | n | n | T | T |
| 335 | L | | | | | n | n | T | T |
| 336 | E | | | | | n | n | T | T |
| 337 | L | | | | | n | n | T | T |
| 338 | T | | | | | n | n | T | T |
| 339 | A | | | | | n | n | T | T |
| 340 | E | | | | | n | n | T | T |
| 341 | E | | | | | n | n | T | T |
| 342 | W | 8.32 | 120.0 | | T338 | y | y | T | T |
| 343 | K | 7.97 | 120.2 | | A339 | y | y | T | T |
| 344 | K | 8.92 | 122.2 | | E340 | y | y | T | T |
| 345 | K | 8.89 | 119.4 | | E341 | y | y | T | T |
| 346 | Y | | | ovlp | W342 | | y | | T |
| 347 | E | 8.78 | 121.8 | | K343 | y | y | T | T |
| 348 | K | 7.98 | 120.1 | | K344 | y | y | T | T |
| 349 | E | 8.37 | 122.6 | | K345 | y | y | T | T |
| 350 | K | 7.89 | 124.4 | | Y346 | y | y | T | T |
| 351 | E | | | weak | E347 | n | n | FN | FN |
| 352 | K | 8.10 | 122.0 | | K348 | y | y | T | T |
| 353 | N | 8.42 | 120.0 | | E349 | y | n | T | FN |
| 354 | K | 8.05 | 119.7 | | K350 | y | y | T | T |
| 355 | A | | | | E351 | n | n | FN | FN |
| 356 | L | 8.63 | 119.8 | | K352 | y | y | T | T |
| 357 | K | 8.55 | 119.0 | | N353 | y | y | T | T |
| 358 | S | | | | K354 | n | n | FN | FN |
| 359 | V | 8.30 | 119.9 | | A355 | y | y | T | T |
| 360 | I | 8.50 | 121.1 | | L356 | y | n | T | FN |
| 361 | Q | 7.87 | 117.7 | | K357 | y | y | T | T |
| 362 | H | 8.02 | 118.0 | | S358 | y | n | T | FN |
| 363 | L | 8.79 | 121.5 | | V359 | y | n | T | FN |

| | | | | | | | | | |
|---|---|---|---|---|---|---|---|---|---|
| 364 | E | 9.00 | 118.4 | | I360 | y | y | T | T |
| 365 | V | 7.92 | 121.1 | | Q361 | y | y | T | T |
| 366 | E | 8.04 | 117.3 | | H362 | y | y | T | T |
| 367 | L | 9.08 | 120.1 | | L363 | y | y | T | T |
| 368 | N | 8.34 | 116.9 | | E364 | y | y | T | T |
| 369 | R | 7.76 | 120.2 | | E366 | y | n | T | FN |
| 370 | W | | | | | n | n | T | T |
| 371 | R | | | | | n | n | T | T |
| 372 | N | | | | | n | n | T | T |
| 373 | G | | | | | n | n | T | T |
| 374 | E | | | | | n | n | T | T |
| 375 | A | | | | | n | n | T | T |
| 376 | V | | | | | n | n | T | T |

[a] For table legend please see Table S1

**Table S3 CspA: H-bonding and surviving $^{1}H$-$^{15}N$ HSQC correlations at pH 10.6.**[a]

| # | aa | $H^N$ | N | notes | HB | hi-pH | HX[a] | HX TMAO[b] | T/F hi-pH | T/F HX | T/F TMAO[b] |
|---|---|---|---|---|---|---|---|---|---|---|---|
| 1 | M | | | | | n | n | n | T | T | T |
| 2 | S | | | | | n | n | n | T | T | T |
| 3 | G | | | | | n | n | n | T | T | T |
| 4 | K | 8.13 | 121.8 | ovlp I8 | | | n | y | | T | FP |
| 5 | M | 8.52 | 125.7 | | F53 | y | n | y | T | FN | T |
| 6 | T | 7.56 | 110.3 | ovlp G26 | | | n | y | | T | FP |
| 7 | G | 8.86 | 111.6 | | V51 | y | y | y | T | T | T |
| 8 | I | 8.13 | 121.8 | ovlp K4 | T22 | | y | y | | T | T |
| 9 | V | 8.74 | 129.3 | | Q49 | y | y | y | T | T | T |
| 10 | K | 9.04 | 139.0 | | F20 | y | y | y | T | T | T |
| 11 | W | 7.33 | 114.0 | | F20 | y | y | y | T | T | T |
| 12 | F | | | | | n | n | n | T | T | T |
| 13 | N | 7.93 | 129.6 | | F18 | y | n | y | T | FN | T |
| 14 | A | | | | | n | n | n | T | T | T |
| 15 | D | | | | | n | n | n | T | T | T |
| 16 | K | | | ovlp S35 | | | | | | | |
| 17 | G | 8.05 | 110.1 | | N13 | y | y | y | T | T | T |
| 18 | F | 6.55 | 115.0 | | N13 | y | y | y | T | T | T |
| 19 | G | 7.93 | 108.0 | | V32 | y | y | y | T | T | T |
| 20 | F | 7.82 | 117.7 | | W11 | y | y | y | T | T | T |
| 21 | I | 9.33 | 124.4 | | V30 | y | y | y | T | T | T |
| 22 | T | 9.26 | 129.6 | | I8 | y | y | y | T | T | T |
| 23 | P | | | | | | | | | | |
| 24 | D | 9.11 | 126.1 | | T6 | y | y | y | T | T | T |
| 25 | D | | | | | n | n | n | T | T | T |
| 26 | G | 7.56 | 110.3 | ovlp T6 | P23 | | n | y | | FN | T |
| 27 | S | | | | | n | n | n | T | T | T |
| 28 | K | | | | | n | n | y | T | T | FP |
| 29 | D | 7.81 | 121.0 | | | y | y | y | FP | FP | FP |
| 30 | V | 9.21 | 123.7 | | I21 | y | y | y | T | T | T |
| 31 | F | 8.48 | 132.9 | | P62 | y | y | y | T | T | T |
| 32 | V | 7.68 | 127.2 | | G19 | y | y | y | T | T | T |
| 33 | H | 8.69 | 130.5 | | S35 OG | y | n | n | T | T | T |
| 34 | F | 8.18 | 126.0 | | G17 | y | n | y | T | T | T |
| 35 | S | | | ovlp K16 | | | | | | | |
| 36 | A | | | | H33 | n | n | n | FN | FN | FN |
| 37 | I | 7.35 | 122.2 | | F34 | y | n | n | T | FN | FN |
| 38 | Q | 8.91 | 130.2 | | V67 | y | | | T | | |
| 39 | N | | | | | n | n | n | T | T | T |
| 40 | D | | | | | n | n | n | T | T | T |

| | | | | | | | | | | | |
|---|---|---|---|---|---|---|---|---|---|---|---|
| 41 | G | | | | | n | n | n | T | T | T |
| 42 | Y | | | | | n | n | n | T | T | T |
| 43 | K | | | | | n | n | n | T | T | T |
| 44 | S | | | | | n | n | n | T | T | T |
| 45 | L | 5.76 | 121.9 | | | y | n | y | FP | T | FP |
| 46 | D | 8.08 | 122.2 | | Q41 OE1 | y | n | n | T | FN | FN |
| 47 | E | 8.63 | 124.0 | ovlp I55 | | | n | y | | T | FP |
| 48 | G | 9.00 | 116.7 | | V9 | y | y | y | T | T | T |
| 49 | Q | 7.65 | 123.6 | | D46 | y | y | y | T | T | T |
| 50 | K | 8.69 | 129.1 | | | y | y | y | FP | FP | FP |
| 51 | V | 8.50 | 118.5 | | G7 | y | y | y | T | T | T |
| 52 | S | 9.04 | 117.7 | | T68 | y | y | y | T | T | T |
| 53 | F | 8.56 | 120.0 | | M5 | y | y | y | T | T | T |
| 54 | T | 8.87 | 113.3 | | G65 | y | y | y | T | T | T |
| 55 | I | 8.63 | 124.0 | ovlp E47 | | | n | y | T | T | FP |
| 56 | E | 9.10 | 130.6 | | A63 | y | n | y | T | FN | T |
| 57 | S | | | | | n | n | n | T | T | T |
| 58 | G | | | | | n | n | n | T | T | T |
| 59 | A | | | | | n | n | n | T | T | T |
| 60 | K | | | | | n | n | n | T | T | T |
| 61 | G | | | | | n | n | n | T | T | T |
| 62 | P | | | | | n | n | | T | T | |
| 63 | A | 8.68 | 128.4 | | E56 | y | n | y | T | FN | T |
| 64 | A | 8.81 | 125.0 | | F31 | y | n | y | T | FN | T |
| 65 | G | 9.20 | 111.7 | | T54 | y | n | y | T | FN | T |
| 66 | N | | | | | n | n | n | T | T | T |
| 67 | V | 8.04 | 120.8 | | A36 | y | n | y | T | FN | T |
| 68 | T | 8.93 | 123.8 | | S52 | y | n | y | T | FN | T |
| 69 | S | | | | | n | n | n | T | T | T |
| 70 | L | 7.89 | 134.2 | | K50 | y | y | y | T | T | T |

[a] For table legend please see Table S1
[b] HX and T/F TMAO – HX isotope exchange protection in $D_2O$ in the presence of the stabilizing osmolyte TMAO (trimethylamine N-oxide) as described in Jaravine et al (2000) Protein Sci 9: 290. This entry is unique to the CspA protein.

**Table S4 LysN: H-bonding and surviving $^{1}H$-$^{15}N$ HSQC correlations at pH 11.0.**[a]

| # | aa | $H^N$ | N | notes | HB | hi-pH | HX[a] | T/F hi-pH | T/F HX |
|---|---|---|---|---|---|---|---|---|---|
| 32 | Q | | | | | n | n | T | T |
| 33 | G | | | | | n | n | T | T |
| 34 | I | | | | | n | n | T | T |
| 35 | A | | | | | n | n | T | T |
| 36 | F | | | | | n | n | T | T |
| 37 | P | | | | | n | n | T | T |
| 38 | N | | | | | n | n | T | T |
| 39 | D | | | | | n | n | T | T |
| 40 | F | | | | | n | n | T | T |
| 41 | R | | | | | n | n | T | T |
| 42 | R | | | | | n | n | T | T |
| 43 | D | | | | | n | n | T | T |
| 44 | H | 8.31 | 114.7 | weak | A69 | y | n | T | FN |
| 45 | T | | | | | n | n | T | T |
| 46 | S | | | | | n | n | T | T |
| 47 | D | 8.88 | 118.5 | ovlp | | | | | |
| 48 | Q | | | | | n | y | T | FP |
| 49 | L | | | | T45 | n | n | FN | FN |
| 50 | H | | | | | n | y | T | FP |
| 51 | A | | | weak | D47 | y | y | T | T |
| 52 | E | 7.70 | 111.9 | | Q48 | y | y | T | T |
| 53 | F | 8.11 | 110.7 | | L49 | y | y | T | T |
| 54 | D | | | ovlp | | | | | |
| 55 | G | | | | | n | n | T | T |
| 56 | K | 7.45 | 120.0 | weak | F53 | y | n | T | FN |
| 57 | E | 9.13 | 118.8 | | E50 OE2 | y | n | T | FN |
| 58 | N | | | | | n | n | T | T |
| 59 | E | | | | | n | n | T | T |
| 60 | E | | | | | n | n | T | T |
| 61 | L | 8.12 | 116.4 | | E57 | y | y | T | T |
| 62 | E | 8.96 | 118.5 | | N58 | y | y | T | T |
| 63 | A | | | | | n | n | T | T |
| 64 | L | 7.72 | 115.4 | | E60 | y | y | T | T |
| 65 | N | 6.83 | 110.4 | | L61 | y | y | T | T |
| 66 | I | | | | | n | n | T | T |
| 67 | E | | | | | n | n | T | T |
| 68 | V | 8.92 | 114.9 | | G126 | y | y | T | T |
| 69 | A | 9.48 | 122.7 | weak | D43 OD1 | y | y | T | T |
| 70 | V | 9.12 | 111.1 | | A124 | y | y | T | T |
| 71 | A | | | | | n | y | T | FP |

| | | | | | | | | | |
|---|---|---|---|---|---|---|---|---|---|
| 72 | G | 9.08 | 126.0 | | L122 | y | y | T | T |
| 73 | R | 8.68 | 117.2 | N/A | G119 | | y | | T |
| 74 | M | 9.27 | 122.4 | | D120 | y | y | T | T |
| 75 | M | 9.08 | 126.0 | | T87 | y | y | T | T |
| 76 | T | 7.55 | 105.6 | | | y | y | FP | FP |
| 77 | R | | | | | n | n | T | T |
| 78 | R | 9.21 | 125.8 | | F85 | y | n | T | FN |
| 79 | I | | | ovlp | | | | | |
| 80 | M | 9.24 | 128.3 | | A83 | y | n | T | FN |
| 81 | G | | | | | n | n | T | T |
| 82 | K | | | | | n | n | T | T |
| 83 | A | | | | | n | n | T | T |
| 84 | S | 9.08 | 110.2 | | V99 | y | y | T | T |
| 85 | F | 8.78 | 117.7 | | R87 | y | y | T | T |
| 86 | V | 9.26 | 121.8 | | L97 | y | y | T | T |
| 87 | T | 8.19 | 118.8 | | T76 | y | y | T | T |
| 88 | L | 9.12 | 126.7 | | I95 | y | y | T | T |
| 89 | Q | 9.07 | 120.9 | | R73 | y | y | T | T |
| 90 | D | 9.63 | 128.8 | weak | G93 | y | y | T | T |
| 91 | V | | | | | n | n | T | T |
| 92 | G | | | | | n | n | T | T |
| 93 | G | 7.72 | 107.7 | weak | D90 | y | n | T | FN |
| 94 | R | | | | | n | n | T | T |
| 95 | I | 8.99 | 116.7 | | L88 | y | y | T | T |
| 96 | Q | 7.60 | 123.3 | | T87 OG1 | y | y | T | T |
| 97 | L | 9.94 | 119.8 | | V86 | y | y | T | T |
| 98 | Y | 9.60 | 122.1 | | I138 | y | y | T | T |
| 99 | V | 8.42 | 127.7 | | S84 | y | y | T | T |
| 100 | A | | | | | n | n | T | T |
| 101 | R | 7.42 | 120.6 | | K82 | y | n | T | FN |
| 102 | D | | | | | n | n | T | T |
| 103 | D | 7.42 | 115.3 | | A100 | y | y | T | T |
| 104 | L | 6.89 | 116.6 | | A100 | y | y | T | T |
| 105 | P | | | | | | | | |
| 106 | E | | | | | n | n | T | T |
| 107 | G | | | | | n | n | T | T |
| 108 | V | 7.81 | 117.7 | | P105 | y | y | T | T |
| 109 | Y | | | | | n | n | T | T |
| 110 | N | | | | | n | n | T | T |
| 111 | E | | | | G107 | n | n | FN | FN |
| 112 | Q | | | N/A | | | | | |
| 113 | F | 7.07 | 112.1 | | V108 | y | y | T | T |
| 114 | K | 7.74 | 115.9 | | Y109 | y | y | T | T |
| 115 | K | 7.09 | 112.9 | weak | Q112 | y | y | T | T |
| 116 | W | 6.89 | 120.8 | | F113 | y | y | T | T |
| 117 | D | 9.40 | 119.5 | | D120 OD2 | y | n | T | FN |

| | | | | | | | | | |
|---|---|---|---|---|---|---|---|---|---|
| 118 | L | | | | | n | n | T | T |
| 119 | G | 9.30 | 110.2 | | M74 | y | y | T | T |
| 120 | D | 8.14 | 119.5 | | D117 | y | y | T | T |
| 121 | I | 8.52 | 117.1 | | T147 | y | y | T | T |
| 122 | L | 8.96 | 124.8 | | G72 | y | y | T | T |
| 123 | G | 8.84 | 103.6 | | R144 | y | y | T | T |
| 124 | A | 9.09 | 124.5 | | V70 | y | y | T | T |
| 125 | K | 8.44 | 117.6 | | E142 | y | y | T | T |
| 126 | G | 8.52 | 106.4 | | V68 | y | y | T | T |
| 127 | K | | | | | n | y | T | FP |
| 128 | L | 8.41 | 121.8 | | I66 | y | y | T | T |
| 129 | F | | | | S137 | n | y | FN | T |
| 130 | K | 8.92 | 116.4 | | N58 OD1 | y | y | T | T |
| 131 | T | | | | E135 | n | n | FN | FN |
| 132 | K | 7.79 | 108.8 | ovlp | | | | | |
| 133 | T | | | | | n | n | T | T |
| 134 | G | | | | | n | n | T | T |
| 135 | E | | | N/A | | | | | |
| 136 | L | | | | | n | n | T | T |
| 137 | S | 9.02 | 119.0 | weak | F129 | y | y | T | T |
| 138 | I | 8.51 | 117.3 | | Q96 | y | y | T | T |
| 139 | H | 10.36 | 128.8 | | K127 | y | y | T | T |
| 140 | C | 8.46 | 122.9 | | Y98 | y | y | T | T |
| 141 | T | 9.67 | 109.7 | | K125 | y | y | T | T |
| 142 | E | 7.96 | 119.1 | | K125 | y | y | T | T |
| 143 | L | | | | | n | n | T | T |
| 144 | R | 9.15 | 120.2 | | G123 | y | y | T | T |
| 145 | L | 8.34 | 122.2 | | | n | n | FP | T |
| 146 | L | 9.01 | 125.7 | | I121 | y | y | T | T |
| 147 | T | | | ovlp | | | | | |
| 148 | K | | | | | n | n | T | T |
| 149 | A | | | | | n | n | T | T |

[a] For table legend please see Table S1

**Table S5 SN: H-bonding and surviving $^{1}H$-$^{15}N$ HSQC correlations at pH 11.0.**[a]

| #FL | aa | $H^N$ | N | notes | HB | hi-pH | HX[a] | T/F hi-pH | T/F HX |
|---|---|---|---|---|---|---|---|---|---|
| 1 | A | | | | | n | n | T | T |
| 2 | T | | | | | n | n | T | T |
| 3 | S | | | | | n | n | T | T |
| 4 | T | | | | | n | n | T | T |
| 5 | K | | | | | n | n | T | T |
| 6 | K | | | | | n | n | T | T |
| 7 | L | | | | | n | n | T | T |
| 8 | H | | | ovlp | | | | | |
| 9 | K | | | | | n | n | T | T |
| 10 | E | 9.26 | 125.7 | | V74 | y | y | T | T |
| 11 | P | | | | | | | | |
| 12 | A | 8.08 | 122.9 | | I72 | y | y | T | T |
| 13 | T | 7.93 | 111.7 | | M26 | y | y | T | T |
| 14 | L | | | | | n | n | T | T |
| 15 | I | | | | | n | y | T | FP |
| 16 | K | 7.97 | 116.6 | | K24 | y | y | T | T |
| 17 | A | | | | | n | n | T | T |
| 18 | I | | | | | n | n | T | T |
| 19 | D | | | | | n | n | T | T |
| 20 | G | | | | | n | n | T | T |
| 21 | D | | | | | n | n | T | T |
| 22 | T | | | | D19 | n | y | FN | T |
| 23 | V | 8.84 | 121.3 | | F34 | y | y | T | T |
| 24 | K | 9.30 | 129.8 | | K16 | y | y | T | T |
| 25 | L | 9.32 | 130.2 | | M32 | y | y | T | T |
| 26 | M | 9.46 | 124.6 | | T13 | y | y | T | T |
| 27 | Y | 8.96 | 132.8 | | Q30 | y | y | T | T |
| 28 | K | 9.32 | 130.2 | | | y | n | FP | T |
| 29 | G | | | | | n | n | T | T |
| 30 | Q | | | ovlp | Y27 | | | | |
| 31 | P | | | | | | | | |
| 32 | M | 9.53 | 128.5 | | L25 | y | y | T | T |
| 33 | T | 8.83 | 125.8 | | | y | n | FP | T |
| 34 | F | 9.51 | 125.4 | | V23 | y | y | T | T |
| 35 | R | 9.57 | 129.1 | | G88 | y | n | T | FN |
| 36 | L | | | | | n | y | T | FP |
| 37 | L | 7.67 | 129.0 | | A90 | y | y | T | T |
| 38 | L | | | | | n | n | T | T |
| 39 | V | 6.93 | 111.7 | | | y | y | FP | FP |
| 40 | D | 8.86 | 124.8 | | K110 | y | n | T | FN |
| 41 | T | | | weak | | | | | |
| 42 | P | | | | | | | | |

| | | | | | | | | | |
|---|---|---|---|---|---|---|---|---|---|
| 43 | E | 8.46 | 124.5 | | E52 OE1 | y | n | T | FN |
| 44 | T | | | weak | D19 OD2 | | | | |
| 45 | K | | | weak | | | | | |
| 46 | H | | | | | n | n | T | T |
| 47 | P | | | | | | | | |
| 48 | K | | | weak | | | | | |
| 49 | K | | | weak | | | | | |
| 50 | G | 8.42 | 113.5 | | H46 | y | n | T | FN |
| 51 | V | | | ovlp | | | | | |
| 52 | E | 8.40 | 125.8 | | E43 | y | n | T | FN |
| 53 | K | | | weak | | | | | |
| 54 | Y | | | weak | | | | | |
| 55 | G | | | | E52 | n | n | FN | FN |
| 56 | P | | | | | | | | |
| 57 | E | | | weak | | | | | |
| 58 | A | 8.74 | 128.9 | | Y54 | y | n | T | FN |
| 59 | S | 7.59 | 114.6 | | G55 | y | n | T | FN |
| 60 | A | 8.30 | 127.1 | | P56 | y | y | T | T |
| 61 | F | 8.10 | 124.7 | | E57 | y | y | T | T |
| 62 | T | 8.58 | 123.1 | | A58 | y | y | T | T |
| 63 | K | 7.54 | 123.1 | | S59 | y | y | T | T |
| 64 | K | 7.84 | 120.3 | | A60 | y | y | T | T |
| 65 | M | 7.65 | 118.8 | | F61 | y | y | T | T |
| 66 | V | 8.09 | 110.5 | | T62 | y | y | T | T |
| 67 | E | 8.64 | 123.9 | | K63 | y | y | T | T |
| 68 | N | 7.46 | 115.8 | | K64 | y | n | T | FN |
| 69 | A | 6.58 | 123.9 | | V66 | y | y | T | T |
| 70 | K | | | | | n | n | T | T |
| 71 | K | | | | D85 OD1 | n | y | FN | T |
| 72 | I | | | | | n | n | T | T |
| 73 | E | 8.68 | 125.9 | | Y93 | y | y | T | T |
| 74 | V | 9.40 | 120.5 | | E10 | y | y | T | T |
| 75 | E | 8.84 | 130.3 | | Y91 | y | y | T | T |
| 76 | F | | | | | n | n | T | T |
| 77 | D | 9.25 | 126.5 | | | y | n | T | FN |
| 78 | K | 10.25 | 121.9 | | D77 OD1 | y | n | T | FN |
| 79 | G | | | | | n | n | T | T |
| 80 | Q | | | | | n | n | T | T |
| 81 | R | | | ovlp | | | | | |
| 82 | T | | | | | n | n | T | T |
| 83 | D | 8.64 | 123.9 | | R87 | y | n | T | FN |
| 84 | K | | | | | n | n | T | T |
| 85 | Y | | | | | n | n | T | T |
| 86 | G | 7.98 | 110.6 | | D83 | n | n | FN | FN |
| 87 | R | | | | | n | n | T | T |
| 88 | G | 8.73 | 111.2 | | T33 | y | n | T | FN |

| | | | | | | | | | |
|---|---|---|---|---|---|---|---|---|---|
| 89 | L | 8.15 | 128.2 | | R81 | y | y | T | T |
| 90 | A | | | | | n | y | T | FP |
| 91 | Y | 9.04 | 124.9 | | E75 | y | y | T | T |
| 92 | I | 7.99 | 125.9 | | N100 OD1 | y | y | T | T |
| 93 | Y | 9.45 | 128.9 | | E73 | y | y | T | T |
| 94 | A | 9.15 | 128.5 | | K87 | y | y | T | T |
| 95 | D | 9.68 | 130.3 | | K71 | y | y | T | T |
| 96 | G | | | | | n | n | T | T |
| 97 | K | | | ovlp | A94 | | y | | T |
| 98 | M | | | | | n | n | T | T |
| 99 | V | 10.16 | 139.0 | | I92 | y | y | T | T |
| 100 | N | 9.67 | 111.3 | | L37 | y | y | T | T |
| 101 | E | 6.05 | 115.1 | | M98 | y | y | T | T |
| 102 | A | 7.84 | 125.9 | | M98 | y | y | T | T |
| 103 | L | 8.14 | 119 | | V99 | y | y | T | T |
| 104 | V | 6.68 | 119.7 | | N100 | y | y | T | T |
| 105 | R | 8.94 | 125.5 | | E101 | y | y | T | T |
| 106 | Q | 7.00 | 114.6 | | A102 | y | y | T | T |
| 107 | G | 7.99 | 109.9 | | V104 | y | y | T | T |
| 108 | L | 7.70 | 117.9 | | L103 | y | y | T | T |
| 109 | A | | | | | n | n | T | T |
| 110 | K | 7.99 | 120.9 | | D40 | y | y | T | T |
| 111 | V | 9.13 | 125.3 | | E129 OE2 | y | n | T | FN |
| 112 | A | 7.83 | 135.1 | | L38 | y | y | T | T |
| 113 | Y | | | | | n | n | T | T |
| 114 | V | 7.75 | 125.4 | | | y | n | FP | T |
| 115 | Y | | | | | n | n | T | T |
| 116 | K | | | | | n | n | T | T |
| 117 | P | | | | | | | | |
| 118 | N | | | | Y115 | n | n | FN | FN |
| 119 | N | 7.86 | 118.9 | | P117 | y | n | T | FN |
| 120 | T | | | | D77 OD2 | n | n | FN | FN |
| 121 | H | 6.72 | 115.9 | | Y91 | y | n | T | FN |
| 122 | E | 7.44 | 123.1 | | N119 | y | n | T | FN |
| 123 | Q | | | ovlp | | | | | |
| 124 | H | | | | | n | n | T | T |
| 125 | L | 7.74 | 122.1 | | H121 | y | n | T | FN |
| 126 | R | 8.82 | 121.0 | | E122 | y | n | T | FN |
| 127 | K | | | ovlp | Q123 | | | | |
| 128 | S | 7.57 | 121.1 | | H124 | y | n | T | FN |
| 129 | E | 8.40 | 127.8 | | L125 | y | y | T | T |
| 130 | A | 7.78 | 122.6 | | R126 | y | y | T | T |
| 131 | Q | 7.43 | 121.3 | | K127 | y | y | T | T |
| 132 | A | 7.92 | 125.6 | | S128 | y | y | T | T |
| 133 | K | 7.99 | 120.5 | | E129 | y | y | T | T |
| 134 | K | 7.79 | 124.4 | | A130 | y | n | T | FN |

| | | | | | | | | | |
|---|---|---|---|---|---|---|---|---|---|
| 135 | E | 7.57 | 119.2 | | Q131 | y | y | T | T |
| 136 | K | 7.75 | 119.7 | | K133 | y | y | T | T |
| 137 | L | 7.55 | 119.8 | | A132 | y | y | T | T |
| 138 | N | | | weak | | | | | |
| 139 | I | | | ovlp | A107 | | | | |
| 140 | W | 7.67 | 122.3 | | A137 | y | n | T | FN |
| 141 | S | | | ovlp | N138 | | | | |
| 142 | E | | | | | n | n | T | T |
| 143 | D | | | | | n | n | T | T |
| 144 | N | | | weak | | | | | |
| 145 | A | | | weak | | | | | |
| 146 | D | | | weak | | | | | |
| 147 | S | | | weak | | | | | |
| 148 | G | | | | | n | n | T | T |
| 149 | Q | | | | | n | n | T | T |

[a] For table legend please see Table S1

**Table S6 GCN4p: H-bonding and surviving $^{1}H$-$^{15}N$ HSQC correlations at pH 9.9.** [a]

| # | aa | $H^N$ | Hα | notes | HB | hi-pH | HX[a] | T/F hi-pH | T/F HX |
|---|---|---|---|---|---|---|---|---|---|
| 1 | G | | | | | n | n | T | T |
| 2 | S | | | | | n | n | T | T |
| 3 | M | | | | | n | n | T | T |
| 4 | K | | | | | n | n | T | T |
| 5 | Q | | | | | n | y | T | FP |
| 6 | L | | | | | n | y | T | FP |
| 7 | E | 8.58 | 123.4 | | M3 | y | y | T | T |
| 8 | D | 8.48 | 122.7 | | K4 | y | n | T | FN |
| 9 | K | 7.84 | 126.2 | | Q5 | y | y | T | T |
| 10 | V | 8.39 | 122.7 | | L6 | y | y | T | T |
| 11 | E | 7.69 | 120.4 | | E7 | y | y | T | T |
| 12 | E | 8.08 | 123.5 | | D8 | y | y | T | T |
| 13 | L | 8.58 | 123.4 | | K9 | y | y | T | T |
| 14 | L | 9.01 | 124.7 | | V10 | y | y | T | T |
| 15 | S | 7.77 | 117.7 | | E11 | y | y | T | T |
| 16 | K | 8.19 | 124.8 | | E12 | y | y | T | T |
| 17 | N | 8.73 | 122.1 | | L13 | y | y | T | T |
| 18 | Y | 8.05 | 123.4 | | L14 | y | y | T | T |
| 19 | H | 7.71 | 121.0 | | S15 | y | y | T | T |
| 20 | L | 8.44 | 123.1 | | K16 | y | y | T | T |
| 21 | E | 8.78 | 122.3 | | N17 | y | y | T | T |
| 22 | N | 7.68 | 121.8 | | Y18 | y | y | T | T |
| 23 | E | 8.11 | 127.1 | | H19 | y | y | T | T |
| 24 | V | 8.64 | 123.0 | | L20 | y | y | T | T |
| 25 | A | 7.66 | 122.6 | | E21 | y | y | T | T |
| 26 | R | 7.90 | 122.4 | | N22 | y | y | T | T |
| 27 | L | 8.40 | 123.1 | | E23 | y | y | T | T |
| 28 | K | 8.92 | 121.6 | | V24 | y | y | T | T |
| 29 | K | 7.24 | 119.8 | | A25 | y | y | T | T |
| 30 | L | 7.38 | 121.5 | | R26 | y | y | T | T |
| 31 | V | 7.64 | 116.7 | | L27 | y | y | T | T |
| 32 | G | 7.73 | 128.6 | | K28 | n | n | FN | FN |
| 33 | E | 8.48 | 122.7 | | | y | n | | |

[a] For table legend please see Table S1

**Table S7 P22iD: H-bonding and surviving $^{1}H$-$^{15}N$ HSQC correlations at pH 10.4.** [a]

| #FL | aa | $H^N$ | N | # | HB | hi-pH | $HX^a$ | T/F hi-pH | T/F HX |
|---|---|---|---|---|---|---|---|---|---|
| 222 | G | | | 1 | | n | n | T | T |
| 223 | S | | | 2 | | n | n | T | T |
| 224 | T | | | 3 | | n | n | T | T |
| 225 | A | | | 4 | | n | n | T | T |
| 226 | T | | | 5 | | n | n | T | T |
| 227 | G | | | 6 | | n | n | T | T |
| 228 | I | | | 7 | | n | n | T | T |
| 229 | T | 8.74 | 116.6 | 8 | S262 OG | y | y | T | T |
| 230 | V | 8.89 | 121.8 | 9 | M338 | y | y | T | T |
| 231 | S | 7.91 | 124.6 | 10 | T260 | y | y | T | T |
| 232 | G | 9.16 | 115.9 | 11 | | y | n | FP | T |
| 233 | A | | | 12 | | n | y | T | FP |
| 234 | Q | 8.22 | 124.3 | 13 | L334 | y | n | T | T |
| 235 | S | | | 14 | | n | n | T | T |
| 236 | F | | | 15 | A314 | n | n | FN | T |
| 237 | K | | | 16 | | n | n | T | T |
| 238 | P | | | 17 | | | | | |
| 239 | V | | | 18 | | n | n | T | T |
| 240 | A | | | 19 | | n | n | T | T |
| 241 | W | | | 20 | | n | n | T | T |
| 242 | Q | | | 21 | | n | n | T | T |
| 243 | L | | | 22 | | n | n | T | T |
| 244 | D | | | 23 | | n | n | T | T |
| 245 | N | | | 24 | | n | n | T | T |
| 246 | D | | | 25 | | n | n | T | T |
| 247 | G | | | 26 | | n | n | T | T |
| 248 | N | | | 27 | | n | n | T | T |
| 249 | K | | | 28 | | n | y | T | FP |
| 250 | V | | | 29 | | n | n | T | T |
| 251 | N | | | 30 | | n | n | T | T |
| 252 | V | | | 31 | | n | n | T | T |
| 253 | D | | | 32 | | n | y | T | FP |
| 254 | N | | | 33 | | n | n | T | T |
| 255 | R | | | 34 | | n | n | T | T |
| 256 | F | | | 35 | | n | y | T | FP |
| 257 | A | 8.63 | 121.7 | 36 | E308 | y | n | T | FN |
| 258 | T | | | 37 | | n | n | T | T |
| 259 | V | 9.30 | 126.6 | 38 | V306 | y | y | T | |
| 260 | T | 9.76 | 123.0 | 39 | | y | y | FP | FP |
| 261 | L | 9.34 | 127.7 | 40 | T304 | y | y | T | |
| 262 | S | 9.48 | 115.8 | 41 | T229 | y | y | T | |
| 263 | A | 7.45 | 119.9 | 42 | | y | y | FP | FP |

| | | | | | | | | | |
|---|---|---|---|---|---|---|---|---|---|
| 264 | T | 7.58 | 102.9 | 43 | G303 | y | y | T | T |
| 265 | T | | | 44 | | n | n | T | T |
| 266 | G | | | 45 | | n | n | T | T |
| 267 | M | 7.95 | 118.7 | 46 | | y | y | FP | FP |
| 268 | K | 9.12 | 120.9 | 47 | D271 OD2 | y | y | T | T |
| 269 | R | | | 48 | | n | n | T | T |
| 270 | G | | | 49 | | n | n | T | T |
| 271 | D | 8.06 | 121.8 | 50 | K268 | y | y | T | T |
| 272 | K | | | 51 | | n | y | T | FP |
| 273 | I | 9.16 | 115.6 | 52 | F295 | y | y | T | T |
| 274 | S | 8.55 | 114.8 | 53 | V340 | y | y | T | T |
| 275 | F | 10.39 | 123.0 | 54 | | y | y | FP | FP |
| 276 | A | 8.29 | 125.1 | 55 | A339 | y | y | T | T |
| 277 | G | | | 56 | | n | n | T | T |
| 278 | V | 8.35 | 124.1 | 57 | F275 | y | y | T | T |
| 279 | K | 9.23 | 127.6 | 58 | | y | y | FP | FP |
| 280 | F | 9.31 | 119.2 | 59 | Q291 | y | n | T | FN |
| 281 | L | | | 60 | | n | n | T | T |
| 282 | G | | | 61 | | n | n | T | T |
| 283 | Q | | | 62 | | n | n | T | T |
| 284 | M | 8.03 | 119.1 | 63 | | y | y | FP | FP |
| 285 | A | | | 64 | | n | n | T | T |
| 286 | K | | | 65 | | n | n | T | T |
| 287 | N | | | 66 | | n | n | T | T |
| 288 | V | | | 67 | | n | n | T | T |
| 289 | L | | | 68 | | n | y | T | FP |
| 290 | A | | | 69 | | n | n | T | T |
| 291 | Q | | | 70 | | n | n | T | T |
| 292 | D | | | 71 | | n | n | T | T |
| 293 | A | | | 72 | V278 | n | n | FN | FN |
| 294 | T | | | 73 | | n | n | T | T |
| 295 | F | 9.46 | 123.0 | 74 | I273 | y | y | T | T |
| 296 | S | 9.04 | 115.8 | 75 | | y | y | FP | FP |
| 297 | V | 9.14 | 124.2 | 76 | D271 | y | y | T | T |
| 298 | V | | | 77 | | n | n | T | T |
| 299 | R | 7.43 | 114.3 | 78 | | y | y | FP | FP |
| 300 | V | | | 79 | | n | y | T | FP |
| 301 | V | 8.32 | 130.6 | 80 | H305 | y | y | T | T |
| 302 | D | | | 81 | T304 | n | y | FN | T |
| 303 | G | | | 82 | | n | n | T | T |
| 304 | T | 8.74 | 108.6 | 83 | | y | n | FP | T |
| 305 | H | | | 84 | | n | y | T | FP |
| 306 | V | 8.77 | 116.4 | 85 | V259 | y | y | T | T |
| 307 | E | 8.92 | 124.7 | 86 | R299 | y | y | T | T |
| 308 | I | 9.45 | 118.3 | 87 | A257 | y | y | T | T |
| 309 | T | 8.59 | 113.5 | 88 | S296 | y | T | T | T |

| | | | | | | | | | |
|---|---|---|---|---|---|---|---|---|---|
| 310 | P | | | 89 | | | | | |
| 311 | K | | | 90 | | n | n | T | T |
| 312 | P | | | 91 | | | | | |
| 313 | V | 9.70 | 130.3 | 92 | K311 | y | y | T | T |
| 314 | A | 9.04 | 131.6 | 93 | | y | y | FP | FP |
| 315 | L | 8.71 | 121.9 | 94 | V330 | y | y | T | T |
| 316 | D | 7.35 | 113.6 | 95 | | y | y | FP | FP |
| 317 | D | 7.10 | 118.0 | 96 | | y | n | FP | T |
| 318 | V | | | 97 | | n | y | T | FP |
| 319 | S | | | 98 | | n | y | T | FP |
| 320 | L | | | 99 | | n | n | T | T |
| 321 | S | | | 100 | | n | n | T | T |
| 322 | P | | | 101 | | | | | |
| 323 | E | 8.77 | 116.4 | 102 | | y | y | FP | T |
| 324 | Q | 7.76 | 117.7 | 103 | S99 | y | y | T | T |
| 325 | R | 8.57 | 119.6 | 104 | S321 | y | n | T | FN |
| 326 | A | | | 105 | | n | n | T | T |
| 327 | Y | 7.68 | 115.7 | 106 | E324 | y | n | T | FN |
| 328 | A | 7.61 | 122.0 | 107 | R325 | y | y | T | T |
| 329 | N | 9.87 | 118.0 | 108 | V313 | y | y | T | T |
| 330 | V | | | 109 | | n | y | T | FP |
| 331 | N | 9.52 | 119.0 | 110 | K279 | y | y | T | T |
| 332 | T | 7.55 | 115.2 | 111 | | y | y | FP | FP |
| 333 | S | 6.90 | 112.6 | 112 | | y | y | FP | FP |
| 334 | L | 8.11 | 117.2 | 113 | E234 | y | y | T | T |
| 335 | A | 8.26 | 130.5 | 114 | | y | y | FP | FP |
| 336 | D | 8.34 | 117.8 | 115 | A233 | y | y | T | T |
| 337 | A | 8.69 | 123.2 | 116 | A335 | y | y | T | T |
| 338 | M | 7.65 | 120.3 | 117 | | y | y | FP | FP |
| 339 | A | 8.67 | 126.8 | 118 | | y | y | FP | FP |
| 340 | V | 7.76 | 121.2 | 119 | I228 | y | y | T | T |
| 341 | N | 8.92 | 126.3 | 120 | S274 | y | y | T | T |
| 342 | I | 8.25 | 122.7 | 121 | | y | n | FP | T |
| 343 | L | 8.96 | 129.6 | 122 | K272 | y | n | T | T |
| 344 | N | | | 123 | | n | n | T | T |
| 345 | V | | | 124 | | n | n | T | T |

[a] For table legend please see Table S1

**Table S8 Cus3iD: H-bonding and surviving $^{1}H$-$^{15}N$ HSQC correlations at pH 10.3.** [a]

| #FL | aa | $H^N$ | N | # | HB | hi-pH | HX[a] | T/F hi-pH | T/F HX |
|---|---|---|---|---|---|---|---|---|---|
| 223 | G | | | 1 | | n | n | T | T |
| 224 | S | | | 2 | | n | n | T | T |
| 225 | T | | | 3 | | n | n | T | T |
| 226 | E | 8.39 | 123.0 | 4 | S224 OG | y | y | T | T |
| 227 | S | | | 5 | | n | n | T | T |
| 228 | L | | | 6 | | n | n | T | T |
| 229 | T | 8.82 | 114.3 | 7 | S258 OG | y | y | T | T |
| 230 | V | 8.73 | 117.9 | 8 | A330 | y | y | T | T |
| 231 | S | 8.69 | 125.3 | 9 | T256 | y | n | T | FN |
| 232 | G | | | 10 | | n | n | T | T |
| 233 | Q | | | 11 | | n | n | T | T |
| 234 | P | | | 12 | | | | | |
| 235 | E | | | 13 | | n | n | T | T |
| 236 | H | 8.85 | 124.4 | 14 | I306 | y | n | T | FN |
| 237 | K | 8.49 | 114.1 | 15 | | y | n | FP | T |
| 238 | V | 8.73 | 120.3 | 16 | S316 OG | y | y | T | T |
| 239 | E | | | 17 | | n | n | T | T |
| 240 | A | | | 18 | | n | n | T | T |
| 241 | K | | | 19 | | n | n | T | T |
| 242 | D | | | 20 | M246 | | n | | FN |
| 243 | S | | | 21 | | n | n | T | T |
| 244 | N | | | 22 | | n | n | T | T |
| 245 | G | | | 23 | D242 | n | n | FN | FN |
| 246 | M | | | 24 | | n | n | T | T |
| 247 | P | | | 25 | | | | | |
| 248 | V | 7.88 | 123.1 | 26 | A240 | | n | | FN |
| 249 | D | | | 27 | | n | n | T | T |
| 250 | N | | | 28 | | n | n | T | T |
| 251 | R | | | 29 | | n | n | T | T |
| 252 | Q | | | 30 | | n | n | T | T |
| 253 | G | | | 31 | I302 | n | n | FN | FN |
| 254 | T | 8.40 | 111.5 | 32 | | n | y | T | FP |
| 255 | I | 9.36 | 118.2 | 33 | V300 | y | y | T | T |
| 256 | T | 9.96 | 119.4 | 34 | | y | y | FP | FP |
| 257 | V | 8.63 | 117.5 | 35 | Q298 | y | y | T | T |
| 258 | S | 8.76 | 112.8 | 36 | T229 | y | n | T | FN |
| 259 | A | 7.38 | 121.3 | 37 | A258 OG | y | n | T | FN |
| 260 | S | | | 38 | | n | n | T | T |
| 261 | G | | | 39 | | n | n | T | T |
| 262 | L | | | 40 | | n | n | T | T |
| 263 | Q | 9.15 | 119.5 | 41 | D266 OD2 | y | y | T | T |

| | | | | | | | | | |
|---|---|---|---|---|---|---|---|---|---|
| 264 | V | | | 42 | | n | n | T | T |
| 265 | G | 8.83 | 116.4 | 43 | V292 | y | n | T | FN |
| 266 | D | | | 44 | Q263 | | y | | T |
| 267 | A | 8.93 | 123.1 | 45 | N366 OD1 | y | y | T | T |
| 268 | F | 9.47 | 117.8 | 46 | F290 | y | y | T | T |
| 269 | T | 9.21 | 109.0 | 47 | T333 | y | y | T | T |
| 270 | I | 10.98 | 122.7 | 48 | | y | y | FP | FP |
| 271 | A | 8.69 | 130.3 | 49 | A331 | y | y | T | T |
| 272 | G | | | 50 | | n | n | T | T |
| 273 | V | | | 51 | I270 | | y | | T |
| 274 | N | 9.59 | 128.2 | 52 | D323 OD1 | y | y | T | T |
| 275 | S | 8.57 | 113.9 | 53 | Q286 | y | y | T | T |
| 276 | V | 7.19 | 110.7 | 54 | N321 | y | n | T | FN |
| 277 | H | 8.93 | 123.1 | 55 | D282 | y | n | T | FN |
| 278 | Q | | | 56 | | n | n | T | T |
| 279 | I | | | 57 | | n | n | T | T |
| 280 | T | | | 58 | | n | n | T | T |
| 281 | K | | | 59 | | n | n | T | T |
| 282 | D | | | 60 | | n | n | T | T |
| 283 | T | | | 61 | | n | n | T | T |
| 284 | T | | | 62 | | | | | |
| 285 | G | | | 63 | | n | n | T | T |
| 286 | Q | | | 64 | | n | n | T | T |
| 287 | P | | | 65 | | | | | |
| 288 | Q | 8.11 | 124.5 | 66 | V273 | y | y | T | T |
| 289 | V | 8.10 | 127.0 | 67 | | y | y | FP | FP |
| 290 | F | 9.17 | 129.1 | 68 | F268 | y | y | T | T |
| 291 | R | 9.03 | 117.7 | 69 | S303 | y | y | T | T |
| 292 | V | | | 70 | | | | | |
| 293 | L | 9.51 | 130.4 | 71 | T301 | y | y | T | T |
| 294 | A | 7.83 | 119.0 | 72 | | y | y | FP | FP |
| 295 | V | | | 73 | | n | n | T | T |
| 296 | S | | | 74 | | n | n | T | T |
| 297 | G | | | 75 | | n | n | T | T |
| 298 | T | | | 76 | | n | n | T | T |
| 299 | T | 8.35 | 118.5 | 77 | S296 | y | n | T | FN |
| 300 | V | 9.51 | 129.6 | 78 | I255 | y | y | T | T |
| 301 | T | 9.07 | 124.6 | 79 | A294 | y | y | T | T |
| 302 | I | 9.50 | 122.3 | 80 | G253 | y | y | T | T |
| 303 | S | 8.39 | 111.5 | 81 | R291 | y | y | T | T |
| 304 | P | | | 82 | | | | | |
| 305 | K | 7.74 | 109.9 | 83 | | y | y | FP | FP |
| 306 | I | 11.12 | 121.7 | 84 | | y | y | FP | FP |
| 307 | L | 8.44 | 128.5 | 85 | N321 OD1 | y | y | T | T |
| 308 | P | | | 86 | | n | n | T | T |
| 309 | V | 7.79 | 123.0 | 87 | V322 | y | y | T | T |

| | | | | | | | | | |
|---|---|---|---|---|---|---|---|---|---|
| 310 | E | | | 88 | | n | n | T | T |
| 311 | N | | | 89 | | n | n | T | T |
| 312 | T | 9.04 | 126.3 | 90 | E235 OE1 | y | n | T | FN |
| 313 | D | 8.91 | 119.6 | 91 | N311 OD1 | y | y | T | T |
| 314 | V | | | 92 | | n | n | T | T |
| 315 | A | 8.56 | 123.1 | 93 | D313 OD2 | y | y | T | T |
| 316 | S | 7.76 | 110.8 | 94 | D313 | y | y | T | T |
| 317 | R | 8.22 | 125.6 | 95 | R314 | y | y | T | T |
| 318 | P | | | 96 | | | | | |
| 319 | Y | | | 97 | | n | y | T | FP |
| 320 | A | 7.35 | 118.5 | 98 | R317 | y | y | T | T |
| 321 | N | 10.02 | 118.2 | 99 | L307 | y | y | T | T |
| 322 | V | 7.73 | 115.1 | 100 | L307 | y | y | T | T |
| 323 | D | 9.50 | 121.0 | 101 | N274 | y | y | T | T |
| 324 | A | | | 102 | | n | y | T | FP |
| 325 | K | 8.01 | 115.8 | 103 | | y | y | FP | FP |
| 326 | P | | | 104 | | | | | |
| 327 | A | | | 105 | | n | n | T | T |
| 328 | E | | | 106 | | n | n | T | T |
| 329 | S | | | 107 | | n | n | T | T |
| 330 | A | 7.10 | 120.9 | 108 | A327 | y | y | T | T |
| 331 | A | | | 109 | | n | n | T | T |
| 332 | I | 8.32 | 124.3 | 110 | L228 | y | y | T | T |
| 333 | T | 9.00 | 125.2 | 111 | T269 | y | y | T | T |
| 334 | I | | | 112 | | n | n | T | T |
| 335 | L | 9.13 | 129.5 | 113 | A267 | y | y | T | T |
| 336 | N | | | 114 | | n | n | T | T |
| 337 | K | | | 115 | | n | n | T | T |

[a] For table legend please see Table S1

**Table S9 BPTI: H-bonding and surviving HN-Hα TOCSY correlations at pH 10.6.** [a]

| # | aa | $H^N$ | Hα | notes | HB | hi-pH | HX[a] | T/F hi-pH | T/F HX |
|---|---|---|---|---|---|---|---|---|---|
| 1 | R | | | | | n | n | T | T |
| 2 | P | | | | | | | | |
| 3 | D | | | | | n | n | T | T |
| 4 | F | | | | | n | n | T | T |
| 5 | C | 7.58 | 4.34 | | P2 | y | y | T | T |
| 6 | L | 7.55 | 4.50 | | D3 | y | y | T | T |
| 7 | E | 7.51 | 4.58 | | F4 | y | y | T | T |
| 8 | P | | | | | | | | |
| 9 | P | | | | | | | | |
| 10 | Y | 7.85 | 4.86 | | | y | y | FP | FP |
| 11 | T | | | | | n | n | T | T |
| 12 | G | | | weak | | n | n | T | T |
| 13 | A | | | ovlp | | | | | |
| 14 | C | 8.82 | 4.55 | | | y | y | FP | FP |
| 15 | K | | | | | n | n | T | T |
| 16 | A | 8.18 | 4.28 | | G36 | y | y | T | T |
| 17 | R | | | | | n | n | T | T |
| 18 | I | 8.18 | 4.18 | | Y35 | y | y | T | T |
| 19 | I | | | | | n | n | T | T |
| 20 | R | 8.37 | 4.71 | | F33 | y | y | T | T |
| 21 | Y | 9.17 | 5.63 | | F45 | y | y | T | T |
| 22 | F | 9.79 | 5.26 | | Q31 | y | y | T | T |
| 23 | Y | 10.56 | 4.30 | | N43 OD1 | y | y | T | T |
| 24 | N | 7.73 | 4.60 | | L29 | y | y | T | T |
| 25 | A | | | | | n | n | T | T |
| 26 | K | | | | | n | n | T | T |
| 27 | A | | | | | n | n | T | T |
| 28 | G | 8.13 | 3.91,3.65 | | N24 | y | y | T | T |
| 29 | L | 6.81 | 4.61 | | N24 | y | y | T | T |
| 30 | C | | | | | n | n | T | T |
| 31 | Q | 8.79 | 4.82 | | F22 | y | y | T | T |
| 32 | T | 8.04 | 5.39 | | | y | y | FP | FP |
| 33 | F | 9.36 | 4.84 | | R20 | y | y | T | T |
| 34 | V | | | ovlp | | | | | T |
| 35 | Y | 9.47 | 4.86 | | I18 | y | y | T | T |
| 36 | G | 8.61 | 3.26,4.32 | | A11 | y | y | T | T |
| 37 | G | | | | | n | n | T | T |
| 38 | C | 7.70 | 4.94 | | | y | n | FP | T |
| 39 | R | | | | | n | n | T | T |
| 40 | A | | | ovlp | Y35 OH | | n | | T |
| 41 | K | 8.43 | 4.43 | | | y | y | FP | FP |
| 42 | R | | | | | n | n | T | T |

| | | | | | | | | | |
|---|---|---|---|---|---|---|---|---|---|
| 43 | N | | | ovlp | | | n | | T |
| 44 | N | | | ovlp | | | y | | FP |
| 45 | F | 9.95 | 5.11 | | Y21 | y | y | T | T |
| 46 | K | | | | | n | n | T | T |
| 47 | S | | | | | n | n | T | T |
| 48 | A | | | | | n | n | T | T |
| 49 | E | | | | | n | n | T | T |
| 50 | D | | | | | n | n | T | T |
| 51 | C | 7.02 | 1.65 | | S47 | y | y | T | T |
| 52 | M | 8.57 | 4.16 | | A48 | y | y | T | T |
| 53 | R | 8.25 | 3.99 | | E49 | y | y | T | T |
| 54 | T | 7.40 | 4.06 | | D50 | y | y | T | T |
| 55 | C | 8.17 | 4.62 | | C51 | y | y | T | T |
| 56 | G | 7.88 | 3.86 | | L52 | y | y | T | T |
| 57 | G | | | | | n | n | T | T |
| 58 | A | | | | | n | n | T | T |

[a] For table legend please see Table S1

**Table S10 SecA-Ct: H-bonding and surviving HN-Hα TOCSY correlations at pH 10.8.**[a]

| # | aa | $H^N$ | Hα | notes | HB | hi-pH | HX[a] | T/F hi-pH | T/F HX |
|---|---|---|---|---|---|---|---|---|---|
| 1 | G | | | | | n | n | T | T |
| 2 | R | | | | | n | n | T | T |
| 3 | N | 7.64 | 4.80 | | | y | n | FP | T |
| 4 | D | | | | | n | y | T | FP |
| 5 | P | | | | | | | | |
| 6 | C | 7.64 | 4.45 | | | y | n | FP | T |
| 7 | P | | | | | n | n | T | T |
| 8 | C | 9.41 | 4.60 | | P5 | y | y | T | T |
| 9 | G | | | | | n | n | T | T |
| 10 | S | 8.92 | 4.16 | | D4 | y | y | T | T |
| 11 | G | | | | | n | n | T | T |
| 12 | K | 7.83 | 4.45 | | D4 OD1 | y | n | T | FN |
| 13 | K | | | | | n | n | T | T |
| 14 | Y | 9.20 | 4.55 | | D4 OD2 | y | n | T | FN |
| 15 | K | | | | | n | n | T | T |
| 16 | Q | 7.54 | 4.42 | | K12 | y | y | T | T |
| 17 | C | 7.84 | 4.49 | | K13 | y | y | T | T |
| 18 | H | 8.84 | 4.22 | | Y14 | y | y | T | T |
| 19 | G | 8.04 | 3.85 | | Y14 | y | y | T | T |
| 20 | R | | | | | n | n | T | T |
| 21 | L | | | | | n | n | T | T |
| 22 | Q | | | | | n | n | T | T |

[a] For table legend please see Table S1

**Table S11 Sequential high pH unfolding of 5 proteins characterized by pH values above which amide peaks are no longer seen.**

| # | GCN4p pH | Kin-neck pH | P22iD pH | Cus3iD pH | Ubq pH |
|---|---|---|---|---|---|
| 1 | 5.8 | | | | 6.0 |
| 2 | 5.8 | | 6.2 | 6.1 | 6.0 |
| 3 | 6.2 | 6.4 | 7.0 | 6.1 | 13.0 |
| 4 | 8.1 | 6.4 | 7.0 | 13.0 | 13.0 |
| 5 | 8.7 | 6.4 | 7.6 | 7.8 | 13.0 |
| 6 | 9.3 | 6.4 | 7.7 | | 13.0 |
| 7 | 10.2 | 6.4 | 9.6 | 11.2 | 10.9 |
| 8 | 10.2 | 6.4 | 11.8 | 13.0 | 8.8 |
| 9 | 10.2 | 6.4 | 12.8 | 10.3 | 6.0 |
| 10 | 11.4 | 6.4 | 12.8 | 7.8 | 11.9 |
| 11 | 11.4 | 6.4 | 11.2 | 9.2 | 7.0 |
| 12 | 11.4 | 6.4 | 11.8 | | 7.0 |
| 13 | 11.4 | 7.0 | 11.2 | 11.2 | 12.4 |
| 14 | 12.7 | 7.0 | 8.2 | 11.2 | 9.0 |
| 15 | 11.4 | 7.8 | 9.6 | 10.3 | 13.0 |
| 16 | 12.7 | 7.0 | 8.2 | 12.2 | 9.0 |
| 17 | 13.0 | 6.4 | | 9.2 | 13.0 |
| 18 | 12.7 | 7.8 | 7.6 | | 13.0 |
| 19 | 12.7 | 7.8 | 7.0 | 9.2 | |
| 20 | 13.0 | 9.0 | 6.2 | 9.2 | 9.0 |
| 21 | 13.0 | 7.8 | | 9.5 | 13.0 |
| 22 | 12.7 | 11.0 | | 7.8 | 6.0 |
| 23 | 13.0 | 9.0 | | 7.8 | 13.0 |
| 24 | 13.0 | 7.8 | 6.2 | 7.8 | |
| 25 | 13.0 | 10.3 | 7.6 | | 11.4 |
| 26 | 13.0 | 11.0 | 7.7 | 9.2 | 13.0 |
| 27 | 13.0 | 11.0 | 7.6 | 7.8 | 13.0 |
| 28 | 13.0 | 11.0 | 7.7 | | 13.0 |
| 29 | 11.4 | | | 9.5 | 13.0 |
| 30 | 11.4 | 11.0 | 7.6 | | 13.0 |
| 31 | 11.4 | 11.0 | 8.2 | | 10.9 |
| 32 | 8.7 | 11.0 | | 12.2 | 10.0 |
| 33 | | 9.0 | 7.0 | 12.2 | 9.0 |
| 34 | | 11.0 | 7.0 | 11.2 | 10.4 |
| 35 | | 11.0 | 9.6 | 13.0 | 10.4 |
| 36 | | 7.8 | | 10.3 | 10.9 |
| 37 | | 11.0 | 11.8 | 10.3 | |
| 38 | | 11.0 | 12.8 | 7.8 | |
| 39 | | 11.0 | 12.8 | 7.8 | 7.0 |

| | | | | | |
|---|---|---|---|---|---|
| 40 | | 11.0 | 12.8 | 9.5 | 9.0 |
| 41 | | 11.0 | 12.8 | 13.0 | 10.9 |
| 42 | | 11.0 | 11.8 | 9.5 | 12.4 |
| 43 | | 11.0 | 11.2 | 10.3 | 10.4 |
| 44 | | 11.0 | 9.6 | 10.3 | 13.0 |
| 45 | | 11.0 | 7.0 | 12.2 | 13.0 |
| 46 | | 10.3 | 11.8 | 13.0 | 7.0 |
| 47 | | 11.0 | 11.8 | 13.0 | 7.0 |
| 48 | | 9.0 | 8.2 | 11.2 | 10.9 |
| 49 | | 9.0 | | 11.2 | 7.0 |
| 50 | | 9.0 | 12.8 | 7.8 | 13.0 |
| 51 | | 6.4 | 10.4 | 12.2 | |
| 52 | | 7.8 | 12.8 | 13.0 | 9.0 |
| 53 | | 7.8 | 12.8 | 12.2 | 6.0 |
| 54 | | 9.0 | 12.8 | 11.2 | 11.4 |
| 55 | | | 12.8 | | 12.4 |
| 56 | | | 9.6 | 12.2 | 13.0 |
| 57 | | | | 6.1 | 6.0 |
| 58 | | | 12.8 | 6.1 | 10.4 |
| 59 | | | 11.8 | 7.8 | 13.0 |
| 60 | | | 9.6 | 7.8 | 9.0 |
| 61 | | | 6.2 | 7.8 | 12.4 |
| 62 | | | | 6.1 | 9.0 |
| 63 | | | 7.0 | 7.8 | 7.0 |
| 64 | | | 7.0 | 9.2 | 10.9 |
| 65 | | | 7.0 | | 11.9 |
| 66 | | | 7.0 | 12.2 | 10.0 |
| 67 | | | 6.2 | 12.2 | 13.0 |
| 68 | | | 6.2 | 13.0 | 13.0 |
| 69 | | | 6.2 | 13.0 | 7.0 |
| 70 | | | 6.2 | | 12.4 |
| 71 | | | 7.7 | 12.2 | 9.0 |
| 72 | | | 9.6 | 13.0 | |
| 73 | | | 9.6 | 11.2 | 13.0 |
| 74 | | | 12.8 | 6.1 | 7.0 |
| 75 | | | 12.8 | 6.1 | 6.0 |
| 76 | | | 12.8 | 7.8 | 7.0 |
| 77 | | | 12.8 | 10.3 | |
| 78 | | | 12.8 | 13.0 | |
| 79 | | | 12.8 | 13.0 | |
| 80 | | | 11.8 | 13.0 | |
| 81 | | | 9.6 | 9.2 | |
| 82 | | | 6.2 | | |
| 83 | | | 9.6 | 12.2 | |
| 84 | | | 10.4 | 11.2 | |
| 85 | | | 12.8 | 13.0 | |

| | | | | |
|---|---|---|---|---|
| 86 | | | 12.8 | |
| 87 | | | 12.8 | 13.0 |
| 88 | | | 12.8 | 9.5 |
| 89 | | | | 9.5 |
| 90 | | | 10.4 | 9.5 |
| 91 | | | | 12.2 |
| 92 | | | | 9.2 |
| 93 | | | 11.8 | 12.2 |
| 94 | | | 11.8 | 12.2 |
| 95 | | | 10.4 | |
| 96 | | | 10.4 | |
| 97 | | | 8.2 | 12.2 |
| 98 | | | 8.2 | 12.2 |
| 99 | | | 9.6 | 13.0 |
| 100 | | | 8.2 | 13.0 |
| 101 | | | | 13.0 |
| 102 | | | 7.6 | |
| 103 | | | 7.0 | 12.2 |
| 104 | | | 7.7 | |
| 105 | | | 8.2 | |
| 106 | | | 11.2 | 7.8 |
| 107 | | | 11.8 | |
| 108 | | | 11.8 | 11.2 |
| 109 | | | 12.8 | 11.2 |
| 110 | | | 11.8 | 12.2 |
| 111 | | | 12.8 | 13.0 |
| 112 | | | 11.8 | 9.5 |
| 113 | | | 12.8 | 11.2 |
| 114 | | | 12.8 | |
| 115 | | | 11.2 | |
| 116 | | | 11.8 | |
| 117 | | | 12.8 | |
| 118 | | | 11.8 | |
| 119 | | | 12.8 | |
| 120 | | | 12.8 | |
| 121 | | | 8.2 | |
| 122 | | | 9.6 | |
| 123 | | | 7.7 | |
| 124 | | | 9.6 | |
